# Fabrication of Molecular Blocks with High Responsiveness to the Cancer Microenvironment by Ursodeoxycholic Acid

**DOI:** 10.1101/2022.10.17.512468

**Authors:** Kazuki Moroishi, Masahiko Nakamoto, Michiya Matsusaki

## Abstract

In cancer therapy, drug delivery system (DDS) has been widely studied to achieve selective drug accumulation at the tumor site. However, DDS still has a major drawback in that it requires multi-step processes for intracellular delivery, resulting in low efficiency of drug delivery. To overcome this problem, we recently reported a molecular block (MB) that disrupts cancer cell membranes in the cancer microenvironment using deoxycholic acid (DCA). However, the MB showed considerable cytotoxicity even at neutral pH possibly due to the structural hydrophobic property of DCA. Herein, we focused on selecting the most suitable bile acid for an MB that possessed high responsiveness to the cancer microenvironment without cytotoxicity at neutral pH. Cell viabilities of the free bile acids such as DCA, chenodeoxycholic acid (CDCA), cholic acid (CA) and ursodeoxycholic acid (UDCA) were evaluated at neutral pH (pH = 7.4) and a cancer acidic environment (pH = 6.3 ∼ 6.5). The half-maximal inhibition concentration (IC_50_) value of UDCA at pH = 7.4 showed an approximately 7.5-fold higher IC_50_ value than that at pH = 6.3, whereas the other bile acids yielded less than a 4-fold IC_50_ value difference between the same pHs. Biocompatible poly(vinyl alcohol) (PVA) was functionalized with UDCA (PVA-UDCA) for synthesis of higher responsiveness to the cancer microenvironment without cytotoxicity at neutral pH. Importantly, 56% pancreatic cancer cell death was observed at pH = 6.5, whereas only 10% was detected at neutral pH by the PVA-UDCA treatment. However, PVA-DCA indicated almost the same cancer cell death property independent of pH condition. These results suggest PVA-UDCA shows great potential for a new class of MB.

## 1. INTRODUCTION

Anticancer chemotherapies to kill cancer cells by targeting DNA^1,2^ are an effective method in cancer treatment. However, current anticancer drugs have limited tumor selectivity and have difficulty accumulating at the target sites due to the widespread distribution of cancer cells in the body, resulting in severe side effects.^3^ To solve this problem, drug delivery systems (DDS) have been widely studied to improve the selective delivery of drugs to tumors.^4^ Sub-100 nm micelles,^5^ liposomes,^6^ and other nanoparticles^7^ have been used for the accumulation of drugs at the tumor site because of the enhanced permeability and retention (EPR) effect.^8^ However, cancer drug-based DDS strategies often target the nucleus and mitochondria, which requires intracellular delivery through a multi-step process, resulting in insufficient cancer cell death.^9^ Furthermore, their therapeutic efficacy becomes limited due to the acquired drug-resistance of cancer cells following repeated chemotherapy.^10^ Drug-free strategies to overcome these challenges have thus attracted attention. This strategy uses cytotoxic peptide and cationic amphiphile, directly disrupting the cell membrane and killing the cell by self-aggregation^11–13^ or charge reversal^14,15^ in response to the cancer microenvironment. Therefore, membrane disruption strategies are expected not only to achieve low-process delivery of molecules, but also to overcome cell drug-resistance.

We have focused on bile acids as cell membrane disrupting molecules in previous drug-free studies. Bile acids are well known biological surfactants and signaling molecules with diverse physiochemical properties. They are generally biosynthesized from cholesterol in the liver and play roles in various physiological processes such as digestion, glucose regulation, lipid metabolism and antibacterial defense in the small intestine.^16–18^ They are also one of the amphiphilic molecules, causing cell death induction by releasing mixed micelles of surfactants and phosphate lipids by incorporation into the cell membrane,^19^ or cell apoptosis *via* cell death signaling.^20,21^ Recently, the cytotoxicity of hydrophobic bile acids has been exploited in the development of novel cancer drugs, and several bile acid derivatives have shown anti-proliferative activities against a wide variety of human cancer cell lines.^22^

In this context, we have previously reported the development of a molecular block (MB) that recognizes the tumor environment and disrupts cancer cell membranes.^23^ The MB is a 4-arm polyethylene glycol functionalized with deoxycholic acid (DCA) which is highly cytotoxic bile acids that responds to the cancer microenvironment. It is designed to disperse as nanoscale assemblies in the blood for efficient circulation and penetration through the stromal tissues. The DCA in the MB has been reported to induce significant cytotoxicity by aggregation *via* hydrophobic interaction at a micro to millimeter scale in the weak acid condition of the cancer microenvironment.^24^ However, the MB also showed considerable cytotoxicity even at neutral pH. Although it was hypothesized that the pH-responsivity of DCA was insufficient, the contribution of the bile acid properties to the pH-responsive cytotoxicity of MBs has not been investigated.

In this study, we focused on the selection of the most suitable bile acid possessing high responsiveness to the cancer microenvironment for MBs without cytotoxicity at neutral pH. Cell viabilities of the free bile acids such as DCA, chenodeoxycholic acid (CDCA), cholic acid (CA) and ursodeoxycholic acid (UDCA) were evaluated at neutral pH (pH = 7.4) and a cancer acidic environment (pH = 6.3 ∼ 6.5) (Figure 1a). These bile acids are expected to have different cell interaction properties and physical properties such as p*K*a, solubility, and hydrophobicity due to the different position and orientation of the hydroxyl groups.^25,26^

**Figure 1.**
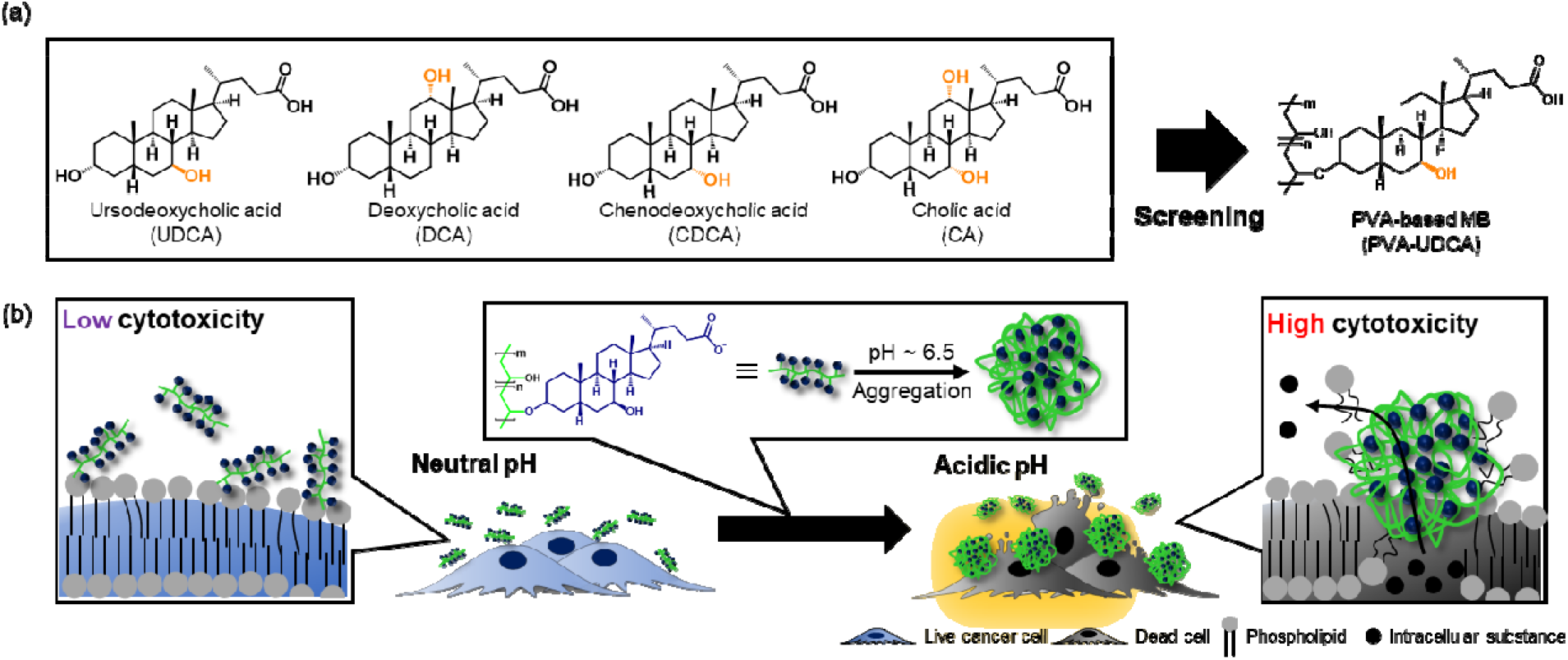
Overview of work. (a) Chemical structure of bile acids and a poly(vinyl) alcohol (PVA)-based molecular block (PVA-UDCA). The UDCA was screened with cytotoxicity evaluation using a cancer cell, and then functionalized with PVA. (b) Schematic illustration of the cytotoxicity mechanism of PVA-UDCA. PVA-UDCA shows low cytotoxicity at neutral pH (pH = 7.4), while high cytotoxicity was observed in an acidic tumor environment (pH = 6.3 ∼ 6.5) by aggregation on the cancer cell surface. Aggregation was caused by protonation of the carboxylate group of bile acids and induced cell death by cell membrane disruption.

The half-maximal inhibition concentration (IC_50_) value of UDCA, which is a known cytoprotective bile acid, showed an approximately 7.5-fold higher IC_50_ value at pH = 7.4 than that at pH = 6.3, whereas the other bile acids indicated less than a 4-fold difference in IC_50_ between the two pHs (DCA: 3.02, CDCA: 4.23, CA: 3.03), suggesting that the UDCA shows highly cancer microenvironment-responsive cell membrane disruption. Based on screening of the bile acids, we developed a MB composed of poly(vinyl alcohol) (PVA) functionalized with UDCA (PVA-UDCA) (Figure 1b). PVA-UDCA showed a higher pH-responsive aggregation property and cytotoxicity than PVA functionalized DCA (PVA-DCA). We revealed that this new class of MB showed highly cancer microenvironment responsive cytotoxicity due to significant changes in hydrophobicity by protonation of UDCA, suggesting this polymer would have further therapeutic potential as a cancer treatment.

## 2. EXPERIMENTAL SECTION

### 2.1. Materials

Deoxycholic acid (DCA), chenodeoxycholic acid (CDCA), ursodeoxycholic acid (UDCA), methanesulfonyl chloride, dehydrated pyridine, 5 M hydrochloric acid (HCl), ethyl acetate, hexane, dimethyl sulfoxide (DMSO), and poly(vinyl) alcohol (PVA) with degree of polymerization 900-1100 were purchased from Wako (Kyoto, Japan). Cholic acid (CA), WST-8 viable cell counting reagent, pyrene and ethanol were purchased from Nacalai Tesque (Kyoto, Japan). Methanol was purchased from Kishida Chemical Co. Ltd. (Osaka, Japan). Phosphate-buffered saline (PBS), and rhodamine B isothiocyanate (TRITC) were purchased from Sigma-Aldrich (St. Louis, USA). Hoechst 33342 was purchased from Invitrogen (Massachusetts, USA). Cellulose tube was purchased from Japan medical science Co. Ltd. (Osaka, Japan)

### 2.2. Cell culture

MiaPaCa-2 was purchased from the American Type Culture Collection (Manassas, VA, USA) and maintained in DMEM supplemented with 10% fetal bovine serum (FBS) and 1% antibiotics. Cells were cultured in 5% CO_2_ and 95% humidified air at 37 °C and passaged every 3 days.

### 2.3. Preparation of medium at weak acidic pH

The medium at weak acidic pH was prepared according to a previously reported method.^27^ A 1 mM HCl solution was added to the medium and incubated for 30 min to allow for equilibrium between the medium and CO_2_, then several μL of HCl and NaOH solution were added every 30 min until the pH of the medium was weakly acidic.

### 2.4. The measurement of critical aggregation concentration (CAC) of bile acids

Transmittance at a wavelength of 450 nm was used for CAC determination.^28^ DCA, CDCA, CA and UDCA were dissolved in PBS at concentrations of 1 ∼ 20 mM and their pH was adjusted to pH = 7.4 and 6.5, respectively. 400 μL of the solutions were used for measurement immediately after preparation. A UV-Vis spectrofluorometer (Nanodrop 2000, Thermo Fisher Scientific, Waltham, US) was set at a fluorescence wavelength ranging from 190 nm to 840 nm. The PBS solution at each pH was used for baseline in this experiment. The CAC value was taken from the intersection of the tangent to the curve at the inflection with the horizontal tangent through the points at low concentrations.

### 2.5. Evaluation of size distribution of bile acids using dynamic light scattering (DLS)

DCA, CDCA, CA and UDCA were dissolved in PBS at a concentration of 2 mM and their pH was adjusted to pH = 7.4 and 6.5, respectively. 500 μL of each solution was used for DLS measurements (Zetasizer Nano-ZS, Malvern, UK) 2 min after preparation.

### 2.6. Evaluation of cytotoxicity of bile acids

The cytotoxicity of bile acids was evaluated by WST-8 assay. MiaPaCa-2 was seeded on a 96-well plate at 1.0×10^4^ cells per well for cytotoxicity measurement at 1, 2, 4, and 6×10^4^ cells per well for standard curve creation using a WST-8 assay. After 15 h incubation, cells for the standard curve were washed with 100 μL of PBS and incubated with 100 μL of 10% WST-8 viable cell counting reagent diluted in phenol red-free DMEM with 1% antibiotics for 2 h at 37 °C and 5% CO_2_. Then, 80 μL of the supernatant was transferred to another 96-well plate and the absorbance at 450 nm was measured using a plate reader (Biotek, VT, US). 10 μM ∼ 10 mM DCA, CDCA, CA and 10 μM ∼ 5 mM UDCA in DMEM medium supplemented with 10% fetal bovine serum (FBS), 1% antibiotics and 0.4% DMSO were prepared at pH = 7.4 or 6.3. The cells were treated with these media for 24 h. Phase contrast microscopy (Ph) images of the cells incubated for 24 h were taken using an EVOS microscope (Thermo Fisher Scientific, Waltham, US). After 24 h, the absorbance of WST-8 was measured using the same procedure as for calibration of the standard curve.

### 2.7. Determination of cell viability and IC_50_ values

Cell viability was determined using the percentage of viable cells by calculating the number of cells in each sample using a standard curve and then using the following formula:

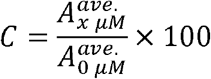

where C is the viable cell rate, 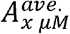 is the average number of living cells cultured with medium containing bile acid x μM for 24 h, and 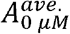 is that of the medium without bile acid. The fitting curve equation used to determine the IC_50_ value is as follows:

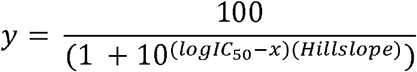

where *y* is the viable cell rate, *x* is the concentration of bile acid and Hillslope is the value calculated by GraphPad Prism.

### 2.8. Mesylation of DCA and UDCA

The mesylation and modification of UDCA or DCA to PVA are depicted in Scheme S1. Mesylation of DCA was performed using previously reported methods.^29^ Briefly, a solution of DCA (**1**) (2.95 g, 7.5 mmol, 1.0 eq) in dehydrated pyridine (50 mL) was stirred at 0 °C in a nitrogen atmosphere. Methanesulfonyl chloride (700 μL 1.2 mmol) was added dropwise, and the solution was stirred at 0 °C for 30 min. The reaction mixture was warmed to room temperature and then stirred for an additional 3 h. To this reaction mixture was added 5 M HCl (100 mL), and the solution was extracted with ethyl acetate (3×100 mL). A solution of mesylated DCA was removed by precipitation using hexane, and dried *in vacuo*. UDCA (**2**) was also mesylated using the same synthetic procedure.

Mesylated DCA (**2**): Yield: 3.95 mg. ^1^H NMR (JEOL, Tokyo, Japan) (400 MHz, Chloroform-*d*): δ = 4.65 (m, 1 H), 4.00 (s, 1 H), 2.99 (s, 3 H, CH_3_SO_3_), 2.44-2.37 (m, 1 H), 2.30-2.23 (m, 1 H), 2.12-1.02 (m, 24 H), 0.99 (d, J = 9.6 Hz, 3 H), 0.92 (s, 3 H), 0.68 (s, 3 H) ppm. IR: ν∼ = 2,925, 2,865, 1,704, 1,344, 914 cm^−1^.

Mesylated UDCA (**5**): Yield: 3.83 mg, ^1^H NMR (400 MHz, Chloroform-*d*): δ = 4.70-4.54 (br. m, 1 H), 3.56 (s, 1 H), 2.99 (s, 3 H, CH_3_SO_3_), 2.38 (m, 1 H), 2.24 (m, 1 H), 2.16-1.03 (m, 24 H), 0.97-0.84 (m, 6 H), 0.66 (s, 3 H) ppm. FT-IR (HORIBA, Kyoto, Japan): ν∼ = 2,927, 2,863, 1,704, 1,336, 923 cm^−1^.

### 2.9. Modification of PVA-DCA and PVA-UDCA

Modification of PVA-DCA and PVA-UDCA followed previously reported methods.^30^ A solution of mesylated DCA (**2**) (1.244 g, 1.7 mmol, 5 eq / -OH) and PVA (15 mg, 3.41×10^−4^ units, 1.0 eq / -OH) in dehydrated DMSO (10 mL) was stirred at 100 °C in a nitrogen atmosphere. Dehydrated pyridine (1.70 mmol, 5 eq / -OH) was added dropwise, and the solution was stirred at 100 °C for 3 days. This reaction mixture was dialyzed in cellulose tubes against methanol, 10 mM NaOH and water for 3 days, 8 h and overnight respectively. The dialyzed mixture was then freeze-dried for 4 days. Mesylated UDCA was also modified using the same synthetic procedure.

PVA-DCA (**3**): Yield: 33 mg, Grafting degree (G.D.): 23.6%. ^1^H NMR (400 MHz, DMSO-*d*_6_): δ = 4.93 (s, 1H), 4.76 (s, 1H), 4.59 (s, 1H), 4.19 (s, 1H), 4.03 (s, 1H), 3.77 (s, 1H), 3.34 (s, 2H, H_2_O), 2.33-0.86 (m, 28H), 0.57 (s, 1H) ppm. IR: ν∼ = 3,384, 2,925, 2,865, 1,704, 1,018 cm^−1^. PVA-UDCA (**6**): Yield: 27 mg, G.D.: 32.5%. ^1^H NMR (400 MHz, DMSO-*d*_6_): δ = 4.93 (s, 1H), 4.76 (s, 1H), 4.59 (s, 1H), 4.01 (s, 1H), 3.72 (s, 1H), 3.35 (s, 2H, H_2_O), 2.32-0.85 (m, 28H), 0.60 (s, 1H) ppm. IR: ν∼ = 3,426, 2,927, 2,863, 1,704, 1,018 cm^−1^.

### 2.10. The measurement of CAC of PVA-DCA and PVA-UDCA

Transmittance at a wavelength of 450 nm was used for CAC determination.^28^ 1 mg of PVA-DCA and PVA-UDCA were dissolved in 50 μL DMSO. These solutions were dissolved in 10 mL PBS at pH = 7.4 or 6.5, respectively, to prepare polymer solutions containing 0.5% DMSO at a concentration of 0.1 mg mL^-1^. Each solution was measured and analyzed using the same procedure as 2.4.

### 2.11. Evaluation of aggregation property of PVA-DCA and PVA-UDCA using Ph microscopy images and DLS measurement

Polymer solutions were prepared in the same manner as 2.10. at pH = 7.4 and 6.5, respectively. The size distributions of each polymer solution at concentrations of 0.1 mg mL^-1^ were measured at 37 °C, 2 min after preparation using DLS measurement. PVA-DCA and PVA-UDCA in PBS at a concentration of 0.1 mg mL^-1^ were observed by Ph microscope using a Countess cell counting slide (Thermo Fisher Scientific, Massachusetts, USA) after 0.5, 3, 6, 12, and 24 h incubation. The number of aggregates of PVA-DCA and PVA-UDCA at pH = 7.4 and 6.5 after 0.5 h was quantified by counting the total number of aggregates of each polymer from these Ph microscope images and normalizing by their area. Size distributions of PVA-DCA and PVA-UDCA aggregates incubated at pH = 7.4 and 6.5 for 0.5, 3, 6, 12, and 24 h were analyzed by measuring the diameter of 100 aggregates each considered as a perfect circle. The analysis of aggregation property using Ph microscopy images was performed with Image J software.

### 2.12. Evaluation of surface charge using measurement of zeta potential

Polymer solutions were prepared in the same manner as 2.10. at pH = 7.4 and 6.5, respectively. The zeta potential of each polymer solution at concentrations of 0.1 mg mL^-1^ were measured at 37 °C, after 0.5, 6, 24 h incubation using Zetasizer Nano-ZS (Malvern, UK).

### 2.13. Concentration-dependent cytotoxicity of PVA-DCA and PVA-UDCA

MiaPaCa-2 was seeded on a 96-well plate at 1.0×10^4^ cells per well for cytotoxicity measurement at 1, 2, 4, and 6×10^4^ cells per well for standard curve creation using WST-8 assay. After 15 h incubation, cells for the standard curve were washed with 100 μL of PBS and incubated with 100 μL of 10% WST-8 viable cell counting reagent diluted in phenol red-free DMEM with 1% antibiotics for 2 h at 37 °C and 5% CO_2_. Then, 80 μL of the supernatant was transferred to another 96-well plate and the absorbance at 450 nm was measured using a plate reader. 0.9 mg of PVA-DCA and PVA-UDCA were dissolved in 15 μL of DMSO. These solutions were dissolved in 3 mL of DMEM supplemented with 10% fetal bovine serum (FBS) and 1% antibiotics at pH = 7.4 or 6.5, to prepare a medium of each polymer at a concentration of 0.3 mg mL^-1^ containing 0.5% DMSO. These were diluted to 0.2, 0.1, 0.05 and 0.01 mg mL^-1^, and then used for cell viability evaluation. MiaPaCa-2 cells were treated with these media at 37 °C and 5% CO_2_ for 24 h. Ph microscopy images of MiaPaCa-2 cells incubated for 24 h were taken using a Ph microscope. After 24 h, the absorbance of WST-8 was measured using the same procedure as for calibration of the standard curve. IC_50_ values of both polymers at pH = 7.4 and 6.5 were determined in the same manner as 2.7.

### 2.14. Contribution of aggregation property of PVA-UDCA to pH-responsive cytotoxicity

MiaPaCa-2 cells were seeded using the same procedure as in 2.13. 1.0 mg of PVA-UDCA was dissolved in 50 μL of DMSO. 10 μL of the PVA-UDCA solution were dissolved in 2 mL of DMEM supplemented with 10% fetal bovine serum (FBS) and 1% antibiotics at pH = 7.4 or 6.5, to prepare a medium of PVA-UDCA at a concentration of 0.1 mg mL^-1^ containing 0.5% DMSO. The medium was diluted to 0.01 mg mL^-1^ and pre-incubated at 37 °C for 24 h to obtain the media containing aggregates of PVA-UDCA. The medium of PVA-UDCA at a concentration of 0.01 mg mL^-1^ pre-incubated for 0 h was also prepared. MiaPaCa-2 cells were treated with the medium pre-incubated at 37 °C and 5% CO_2_ for 0 h and 24 h and cell viability was measured by the same procedure as in 2.13. with a WST-8 assay.

### 2.15. Modification of tetramethyl rhodamine isothiocyanate (TRITC)

Modification of TRITC followed previously reported methods.^31^ The solution of PVA-DCA (2.0 mg, 4.55×10^−5^ units 1.0 eq / -OH), TRITC (1.21 mg, 10.2 μmol, 0.05 eq / -OH), a drop of pyridine and dibutyltin (IV) dilaurate in dehydrated DMSO (2 mL) was stirred at 95 °C for 2 h in a nitrogen atmosphere. This reaction mixture was dialyzed using cellulose tubes against methanol for 6 days and against water for 1 day. The dialyzed mixture was freeze-dried for 7 days. PVA-UDCA was also modified using this procedure. The grafting degree of PVA-DCA and PVA-UDCA was calculated by fluorescence intensity of TRITC using a spectrofluorometer. This was set at an excitation wavelength of 550 nm, fluorescence bandwidth of 2.5 nm, acquisition interval of 0.5 nm, temperature of 25 °C, and fluorescence wavelength of 450 nm to 650 nm. PVA-DCA-TRITC: Yield 10 mg, G.D.: 0.02% PVA-UDCA-TRITC: Yield 4 mg, G.D.: 0.08%.

### 2.15. Evaluation of cell adsorption of PVA-DCA-TRITC and PVA-UDCA-TRITC using confocal images

MiaPaCa-2 cells were seeded on a glass based 96-well plate at 1.0×10^4^ cells per well for cell adsorption measurement. 0.1 mg of PVA-DCA-TRITC and 0.3 mg of PVA-UDCA-TRITC were dissolved in 5 μL and 15 μL of DMSO, respectively, to prepare polymer solutions at a concentration of 20 mg mL^-1^. 1.7 mg of PVA-DCA and PVA-UDCA were also dissolved in 85 μL of DMSO, to prepare polymer solutions at a concentration of 20 mg mL^-1^. The amount of TRITC was adjusted by mixing 12 μL of PVA-DCA-TRITC solution with 18 μL of PVA-DCA solution, or 3 μL of PVA-UDCA-TRITC with 27 μL of PVA-UDCA, respectively. 10 μL of these polymer solutions were dissolved in 2 mL of phenol red-free DMEM supplemented with 10% fetal bovine serum (FBS) and 1% antibiotics at pH = 7.4 or 6.5, to prepare a medium of each polymer at a concentration of 0.1 mg mL^-1^ containing 0.5% DMSO. MiaPaCa-2 cells were treated with these media at pH = 7.4 and 6.5 under 37 °C and 5% CO_2_. Confocal images of MiaPaCa-2 cells incubated for 3 h and 24 h were taken using a confocal laser scanning microscope (FV3000, Olympus, Tokyo, Japan). MiaPaCa-2 incubated for 3 and 24 h were washed 3 times with phenol red-free DMEM adjusted at pH= 7.4 and 6.5, respectively, and treated with the medium diluted 1,000-fold with Hoechst 33342 for 15 min incubation. After washing 3 times, confocal images of MiaPaCa-2 cells incubated for 3 h and 24 h were again taken.

### 2.16. Statistical analysis

The results are expressed as mean ± standard deviation (S.D.). Student’s t-tests were used to test for differences between group means.

## 3. RESULT AND DISCUSSION

### Evaluation of aggregation property and cytotoxicity of bile acids

Bile acids have a rigid hydrophobic steroid core with hydroxyl and carboxylate groups. They can therefore easily form aggregates due to their amphiphilic property.^32,33^ To confirm this aggregation property, we evaluated the critical aggregation concentration (CAC) and size distribution of aggregates of each bile acid in a phosphate-buffered saline (PBS) solution. The CAC, defined as concentration of amphiphilic molecules at which aggregates are formed, was evaluated by an optical transmittance method.^28^ The transmittance at 450 nm of each bile acid solution at pH = 7.4 did not decrease within 20 mM and CAC could not be determined (FigureS1a-d). In contrast, that of DCA, CDCA, and UDCA at pH = 6.5 decreased with increasing concentration, confirming self-aggregation of these bile acids (Figures S1e, f and h). The CAC of DCA at the weak acid condition was determined to be 2.56 mM and that of CDCA was 1.93 mM, which was a slightly higher aggregation property. On the other hand, the CAC of UDCA was 4.18 mM, indicating a lower aggregation property of UDCA. CA did not decrease in transmittance at pH = 6.5 as it did at pH = 7.4 (Figure S1g). In addition, the size distribution of each bile acid at a concentration of 2 mM, which is below the CAC, was measured by dynamic light scattering (DLS) measurements to evaluate nanometer-sized aggregates of bile acids. The size distribution of DCA at pH = 7.4 revealed that aggregates larger than 100 nm were abundantly distributed in the solution. In contrast, that of CDCA, CA and UDCA formed aggregates of less than 2 nm (Figure 2a). At pH = 6.3, the size distribution of UDCA showed that aggregates larger than 100 nm were abundantly distributed. DCA, CDCA and UDCA formed aggregates of larger than 1,000 nm (Figure 2b), suggesting that these bile acids formed large aggregates with a decrease in pH at the nanoscale. However, the size distribution of CA at pH = 6.3 was not markedly different to that at pH = 7.4, indicating no pH-responsive aggregation. CA did not show pH-responsive aggregation properties in the CAC and DLS measurements due to a considerably lower p*K*a (p*K*a = 5.50) than the other bile acids (DCA: 6.30, CDCA: 6.53 and UDCA: 6.25).^25^ This therefore suggests that bile acids with a p*K*a = 6.3 ∼ 6.5 would have a cancer microenvironment-responsive aggregation property.

**Figure 2.**
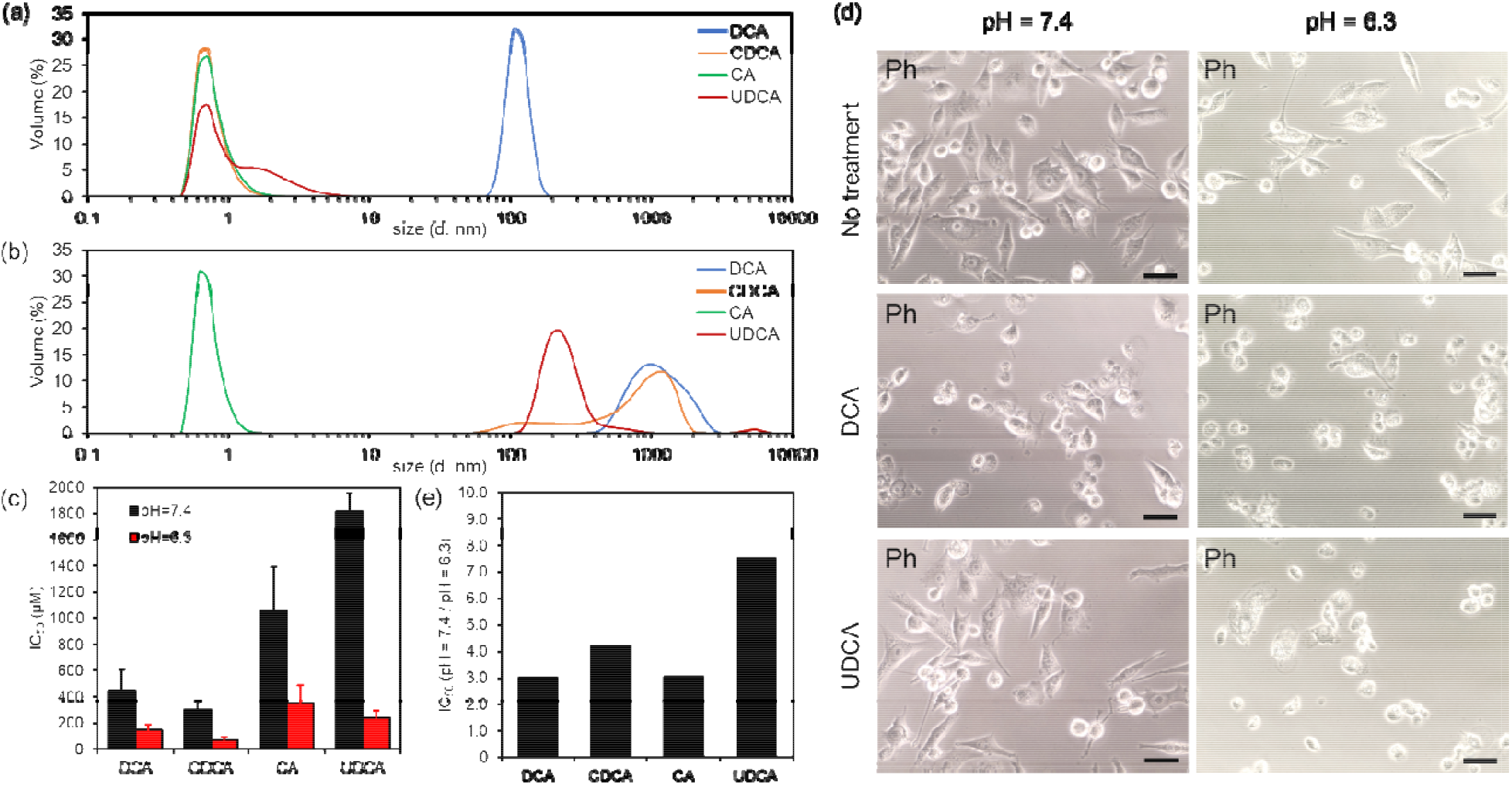
Screening of bile acids. Size distribution of 2 mM DCA (blue), CDCA (orange), CA (green) and UDCA (red) solution at pH = 7.4 (a) and 6.3 (b). n = 6, the data came from technical replicates. (c) The half maximal inhibitory concentration (IC_50_) values of each bile acid using pancreatic (MiaPaCa-2) cells at pH = 7.4 (black) and 6.3 (red) after 24 h incubation. n = 3 biologically independent samples. IC_50_ values were calculated from sigmoidal curve fitting. (d) Phase contrast (Ph) microscopy images of MiaPaCa-2 cells treated with medium (no treatment), 1000 μM DCA and UDCA at pH = 7.4 and 6.3 after 24 h incubation at 37 °C. Scale bars = 50 μm. (e) Ratio of IC_50_ values for pH = 7.4 and pH = 6.3 in DCA, CDCA, CA, and UDCA.

To investigate the cytotoxicity of each bile acid at pH = 7.4 and 6.3, half-maximal inhibition concentration (IC_50_) values were estimated by measuring cell viability of pancreatic cancer (MiaPaCa-2) cells treated with each bile acid after 24 h incubation at pH = 7.4 and 6.3 (Figure S2). The IC_50_ values of DCA and CA at pH = 7.4 were 450 μM and 1,060 μM, respectively, suggesting an approximately 2.4-fold lower cytotoxicity of CA than DCA (Figure 2c). The IC_50_ value of UDCA was 1,820 μM, an approximately 4.0-fold lower cytotoxicity than that of DCA and the lowest value among the four bile acids. On the other hand, the IC_50_ value of CDCA was 300 μM, which was a cytotoxicity about 1.5 times higher than that of DCA and the highest among the four bile acids. These results revealed that IC_50_ values for each bile acid were lower than the concentration at which DLS was measured, suggesting that they could show cytotoxicity before aggregate formation. It has been reported that bile acids can penetrate the hydrophobic region of liposomes in monomeric or submicellar form and interact efficiently with the lipid bilayer membrane.^34,35^ Our results are in good agreement with these reports, indicating that there is no apparent relationship between aggregation and cytotoxicity in the case of bile acids added to cells in monomeric form. In addition, bile acids showed significantly different cytotoxicity at pH = 7.4 due to slight differences in the position of the hydroxyl group. The position of the hydroxyl group would contribute to the differences in hydrophobicity, resulting in different cytotoxicity with each bile acid. It has been reported that hydrophobic bile acids induce cell death *via* apoptosis signaling pathways^20^ and disruption of the cell membrane by releasing mixed micelles of surfactants and phosphate lipids after insertion into the cell membrane.^19,36^ On the other hand, hydrophilic bile acids have cytoprotective activity as inhibit the apoptosis pathway.^21^ The hydrophobicity of bile acids used in this study was reported to be in the order of UDCA < CA < CDCA < DCA in both ionized and protonated states.^37^ Our results revealed that the cytotoxicity of bile acids appear to be closely related to their hydrophobicity. This indicates that DCA and CDCA, which have higher hydrophobicity than CA and UDCA, are high in cytotoxicity probably due to cell membrane disruption or cell death signaling by their hydrophobicity, while CA and UDCA are low in cytotoxicity due to inhibition of cell apoptosis by their hydrophilicity.

Compared to the IC_50_ value of 150 μM for DCA at pH = 6.3, the IC_50_ value for CA was 350 μM, an approximately 2.3-fold lower cytotoxicity and the lowest among the four bile acids at that pH (Figure 2c). The IC_50_ value of UDCA was 240 μM, an approximately 1.6-fold lower cytotoxicity than that of DCA. The IC_50_ value of CDCA was 72 μM, an approximately 2.0 times higher cytotoxicity than DCA and the highest among the four bile acids. These results showed that the IC_50_ values of all bile acids were lower than those at pH = 7.4, suggesting that the increased hydrophobicity due to protonation of each bile acid increases its cytotoxicity. Furthermore, the difference in cytotoxicity of bile acids at pH = 6.3 was smaller than that at pH = 7.4, though the trend at both pHs was similar (Figure 2c). CA with a considerably lower p*K*a than the other bile acids showed the lowest cytotoxicity at pH = 6.3, indicating its moderate hydrophobicity at pH = 6.3 may owing to the abundance of deprotonated carboxylates. In addition, the size of MiaPaCa-2 treated with hydrophobic bile acids such as DCA and CDCA was smaller than that of control cells at both pHs, indicating that excessive hydrophobicity contributed to cell damage regardless of pH (Figures 2d and S3). Conversely, hydrophilic bile acids such as CA and UDCA had little effect on the cell size at pH = 7.4 (Figures 2d and S3). Moreover, the cells treated with UDCA after 24 h incubation showed cell elongation at pH = 7.4 and cell shrink at pH = 6.3, suggesting these bile acids had sharp pH-responsiveness due to hydrophobicity change (Figure 2d). We also evaluated the IC_50_ ratio for pH = 7.4 and 6.3 (Figure 2e). The IC_50_ ratio of UDCA was 7.51, showing the greatest difference in cytotoxicity in response to pH of the four bile acids (DCA: 3.02, CDCA: 4.23 and CA: 3.03).

Taken together, these results suggest that UDCA would be the most suitable bile acid for a cancer therapeutic molecule with hydrophobicity suited to the cancer microenvironment.

### Synthesis and characterization of PVA functionalized with bile acids

The design of a new class of MBs for higher pH-responsive cytotoxicity in a cancer microenvironment, named PVA-DCA and PVA-UDCA was considered using poly(vinyl alcohol) (PVA) as a biocompatible polymer which enables tunable modification of bile acids (Scheme S1). The PVA was modified via mesylation of the bile acids by a previously reported method.^30^ PVA-DCA and PVA-UDCA were successfully synthesized as characterized by FT-IR and ^1^H NMR measurement (Figures S4-S9). The influence of DCA and UDCA unit number on PVA-DCA and PVA-UDCA were evaluated respectively, indicating that the grafting degree (G.D.) of PVA-DCA and PVA-UDCA was 23.6% and 32.5%, respectively.

To confirm whether these polymers have a similar aggregation property to bile acids, we investigated the aggregation property of PVA-DCA and PVA-UDCA in a PBS solution with CAC and evaluated the size of aggregates using DLS measurement and phase contrast (Ph) images (Figure 3a). The CAC of these polymers was evaluated by the same procedure as in the evaluation of the CAC of bile acids. The transmittance of both polymer solutions at 450 nm decreased with an increase in concentration, confirming the self-aggregation of polymers (Figure S10). The CAC values of PVA-DCA and PVA-UDCA at pH = 7.4 were 0.042 mg mL^-1^ and 0.052 mg mL^-1^, respectively, suggesting the lower aggregation property of PVA-UDCA due to the low hydrophobicity of UDCA (Figure 3b). In contrast, the CAC values of PVA-DCA and PVA-UDCA at pH = 6.5 were 0.044 mg mL^-1^ and 0.045 mg mL^-1^, indicating neither polymer changed significantly. These results suggest that the CAC values of PVA-UDCA decrease in response to pH due to increasing hydrophobicity resulting from the protonation of UDCA. On the other hand, the aggregation property of PVA-DCA at pH = 7.4 was comparable to that at pH = 6.5, indicating that PVA-DCA easily forms aggregates even at neutral pH due to its high hydrophobicity.

**Figure 3.**
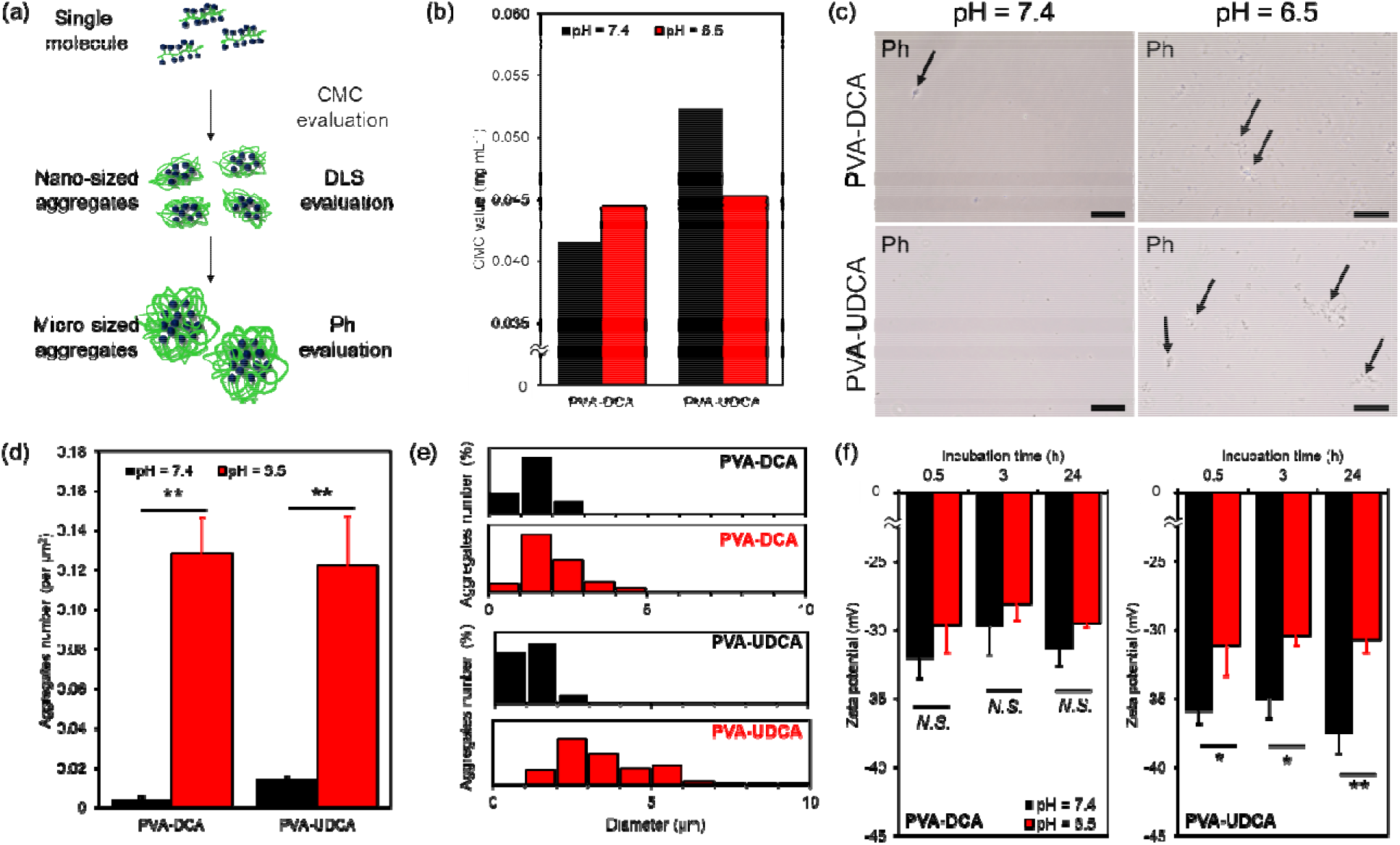
Aggregation property of PVA-DCA and PVA-UDCA in response to a weak acid condition. (a) Procedure of size evaluation. To investigate the aggregation property of PVA-UDCA and PVA-DCA, the CAC of both polymers was analyzed by measuring transmittance at a wavelength of 450 nm. Size distribution of PVA-DCA and PVA-UDCA was measured at 37 °C by DLS at a concentration above CAC. Number and size distribution of micro-sized aggregates were analyzed by phase contrast (Ph) microscopy images. (b) CAC values of PVA-DCA and PVA-UDCA in phosphate buffer saline (PBS) at 37 °C measured by the pyrene method at pH = 7.4 (black) and 6.5 (red). (c) Ph images of 0.1 mg mL^-1^ PVA-DCA and PVA-UDCA at pH = 7.4 and 6.5 after 0.5 h incubation at 37 °C. Scale bars = 50 μm. Black arrows show aggregates of PVA-DCA and PVA-UDCA. (d) Aggregate number of PVA-DCA and PVA-UDCA aggregates at 0.1 mg mL^-1^ in PBS solution at 37 °C incubated for 0.5 h at pH = 7.4 (black) and 6.5 (red). Six areas in the well were randomly selected for the analysis. (e) Size distribution of 0.1 mg mL^-1^ PVA-DCA and PVA-UDCA at pH = 7.4 (black) and pH = 6.5 (red) after 0.5 h incubation at 37 °C. (f) Zeta potential of 0.1 mg mL^-1^ PVA-DCA and PVA-UDCA at pH = 7.4 (black) and pH = 6.5 (red) after 0.5, 3 and 24 h incubation at 37 °C. Statistical analysis was performed using unpaired two-tailed Student’s *t*-test. Data are presented as mean ± S.D. (*N*.*S*.: no significant difference, \**p* < 0.05, \*\**p* < 0.01).

For further investigation, the size of aggregates of polymers at above the CAC was analyzed by DLS measurement and Ph images. The size distribution of PVA-DCA and PVA-UDCA in PBS at 0.1 mg mL^-1^ was measured by DLS to evaluate nanometer-sized aggregates of both polymers. The size distribution of PVA-DCA at pH = 7.4 was comparable to that at pH = 6.5, indicating that PVA-DCA forms large aggregates even at pH = 7.4 due to the high hydrophobicity of the DCA moiety (Figures S11a and b). In contrast, PVA-UDCA formed smaller aggregates than PVA-DCA at pH = 7.4 (Figure S11c), and larger aggregates than PVA-DCA at pH = 6.5 (Figure S11d). These results showed that PVA-UDCA formed large aggregates with a decrease in pH, suggesting high pH-responsiveness for the cancer microenvironment at the nanoscale.

The number of PVA-DCA and PVA-UDCA aggregates at 0.5 h incubation and the size distribution of micrometer-sized aggregates of both polymers at 0.5, 3, 6, 12, and 24 h incubation were evaluated using Ph microscopy images. These images revealed micrometer-sized aggregates of both polymers at pH = 6.5, but few at pH = 7.4 (Figure 3c). Moreover, the number of aggregates of PVA-DCA at a concentration of 0.1 mg mL^-1^ after 0.5 h incubation was 0.0037 per μm^2^ at pH = 7.4 (Figure 3d). In contrast, the number of aggregates at pH = 6.5 was 0.13 per μm^2^, which was about 34-fold higher than that at pH = 7.4. The same conditions for PVA-UDCA showed that the number of aggregates was 0.014 per μm^2^ at pH = 7.4, and approximately 8.6-fold higher at 0.12 per μm^2^ at pH = 6.5. In addition, the size distribution of both polymers at a concentration of 0.1 mg mL^-1^ after 0.5 h incubation at pH = 7.4 revealed smaller than 2 μm aggregates were abundantly distributed in the solution (Figure 3e). In contrast, those of pH = 6.5 showed that formation of aggregates larger than 2 μm was observed in nearly 90% of PVA-UDCA, but only 41% of PVA-DCA, suggesting that PVA-UDCA forms slightly larger aggregates than that of PVA-DCA. Notably, PVA-DCA had the largest number of 1 ∼ 2 μm aggregates at both pHs, indicating the aggregate size did not markedly increase in response to pH. In PVA-UDCA, aggregates of 1 ∼ 2 μm and 2 ∼ 3 μm were most abundant at pH = 7.4 and 6.5, respectively, suggesting the size of PVA-UDCA increased at pH = 6.5 compared to that at pH = 7.4. These results indicate that protonation of the DCA and UDCA moieties would increase hydrophobicity, resulting in a faster increase in aggregate size at pH = 6.5 than at pH = 7.4. The size of aggregates of both polymers at pH = 7.4 and 6.5 markedly increased over time incubation (Figures S12 and S13). Particularly, aggregates of PVA-DCA for 24 h incubation were smaller than that of PVA-UDCA (Figure S12), suggesting that the DCA moiety may have been buried in the aggregates due to high hydrophobicity, limiting polymer interaction.

In addition, the zeta potential of PVA-DCA and PVA-UDCA at pH = 7.4 and 6.5 were evaluated to investigate the surface charge of their aggregates (Figure 3f). The zeta potentials of PVA-DCA at pH = 7.4 and 6.5 after 0.5 h of incubation were −32.1 mV and −29.7 mV, respectively, while these values for PVA-UDCA were −36.0 mV and −31.2 mV, respectively. These results revealed that both polymers at pH = 6.5 showed lower negative charge values than those at pH = 7.4, suggesting anionic charge would decrease by protonation of each bile acid. The zeta potential of PVA-DCA at pH = 7.4 was similar to that at pH = 6.5 at all incubation times, while PVA-UDCA showed significant zeta potential differences between these pHs. DCA has a higher p*K*a value than UDCA, suggesting that DCA has a lower ionization ratio than UDCA at pH = 7.4.^25^ It was suggested that the lower negative charge value of PVA-DCA at both pHs than that of PVA-UDCA is due to a higher p*K*a value than UDCA. In addition, these results may also be due to the DCA moiety being buried in the aggregate and covered by nonionic PVA. The zeta potentials of both polymers did not change over time within 24 hours, suggesting the high stability of the aggregates formed.

Taken together, these results clearly suggested that the number of aggregates of PVA-DCA and PVA-UDCA increased with decreasing pH due to their high hydrophobicity by protonation of the bile acids. However, the aggregate size of PVA-DCA at pH = 7.4 was confirmed to be comparable to that at pH = 6.5 in both nanometer and micrometer size due to its high hydrophobicity. In the case of PVA-UDCA, a marked pH-responsive increase in aggregate size was shown, as the hydrophilicity of PVA-UDCA is higher at pH = 7.4 than that of PVA-DCA. The results of zeta potential showed the difference between PVA-DCA and PVA-UDCA at both pHs, suggesting the difference of aggregation form due to hydrophobic-hydrophobic balance in each polymer.

### Cytotoxicity evaluation of PVA-DCA and PVA-UDCA

The pH-responsive cytotoxicity of PVA-DCA and PVA-UDCA was investigated by treating MiaPaCa-2 with PVA-DCA and PVA-UDCA at different concentrations for 24 h incubation (Figure 4a). The cell viability of MiaPaCa-2 cells treated with PVA-DCA at pH = 7.4 and 6.5 began to decrease at 0.1 mg mL^-1^ and cell attachment was observed even after 24 h incubation at 0.1 mg mL^-1^ by Ph microscopy images (Figure S14a). In contrast, that of PVA-UDCA at pH = 7.4 stayed above 90% up to 0.1 mg mL^-1^, suggesting lower cytotoxicity of PVA-UDCA than that of PVA-DCA. For pH = 6.5, cytotoxicity of PVA-DCA increased from 0.1 mg mL^-1^ as well as pH = 7.4. On the other hand, the cell viability of MiaPaCa-2 cells treated with PVA-UDCA decreased from 0.05 mg mL^-1^, suggesting higher cytotoxicity of PVA-UDCA than PVA-DCA at low concentration. Ph microscopy images of MiaPaCa-2 cells treated with 0.1 mg mL^-1^ of PVA-UDCA after 24 h incubation showed cell elongation at pH = 7.4 and cell detachment at pH = 6.5 (Figure S14b). Importantly, cell viability of MiaPaCa-2 cells treated with 0.1 mg mL^-1^ PVA-DCA was comparable at pH = 6.5 and pH = 7.4 (64% and 72%, respectively) whereas PVA-UDCA showed half the cell viability at pH = 6.5 than at pH = 7.4 (44% and 90%, respectively), indicating that PVA-UDCA would exhibit pH-responsive cytotoxicity. This results also revealed the higher pH-responsive cell viability of PVA-UDCA than that of previous MBs at the same concentration (> 80% at both pHs).^23^ IC_50_ values of both polymers were determined (Figure S15). IC_50_ values of PVA-DCA at pH = 7.4 and 6.5 were 1.16 μM and 1.31 μM, respectively, which were compatible at both pHs. In contrast, IC_50_ values of PVA-UDCA at pH = 7.4 and 6.5 were 1.27 μM and 0.60 μM, respectively, suggesting an approximately 2.1-fold higher cytotoxicity at pH = 6.5 than at pH = 7.4. Importantly, both polymers decreased the IC_50_ values by more than 100-fold at both pH compared to DCA and UDCA alone. This may be due to increased local concentrations of bile acids accumulated in polymers.

**Figure 4.**
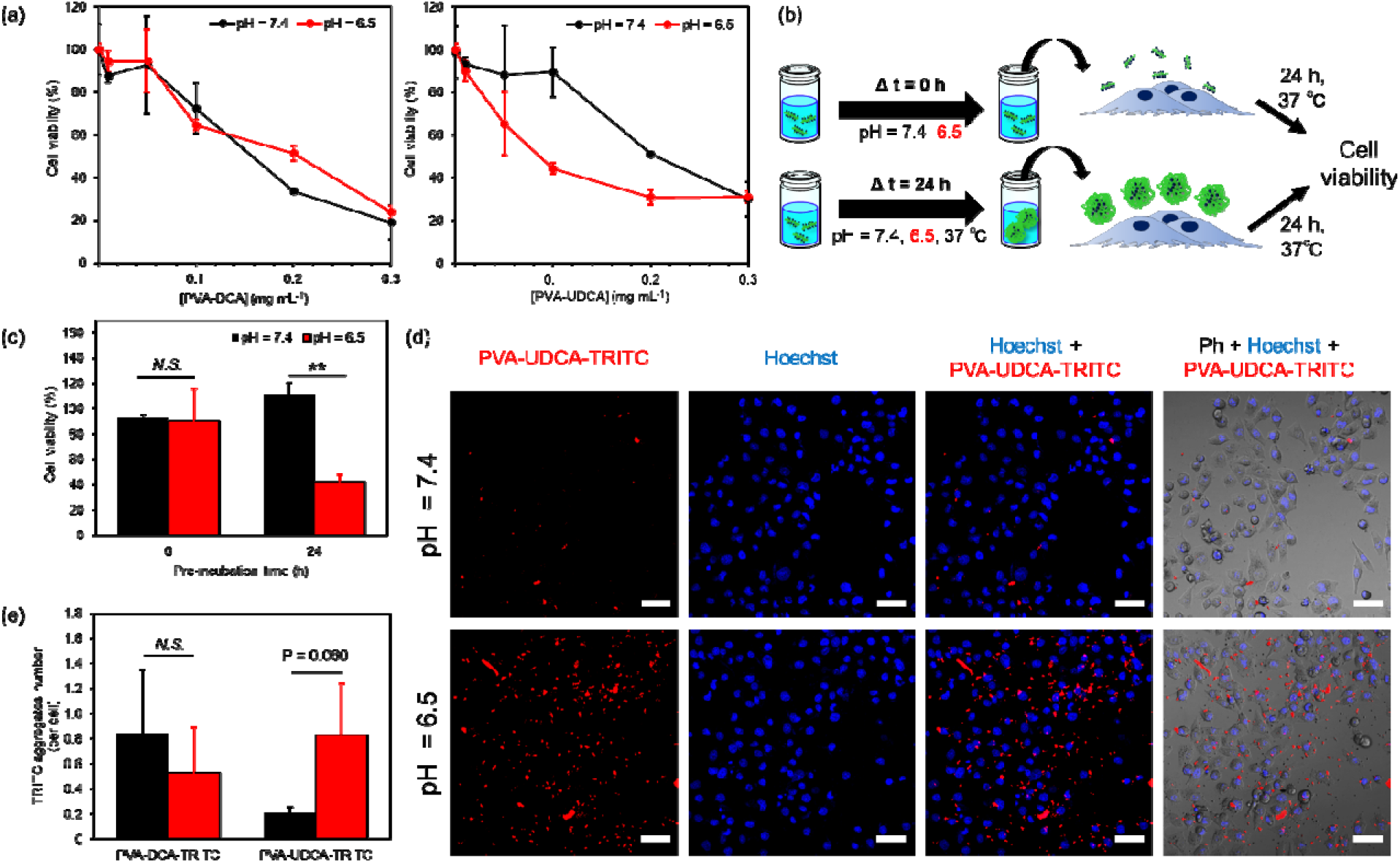
Cytotoxicity effect of PVA-DCA and PVA-UDCA under neutral and weak acid pH condition. (a) Concentration-dependent cytotoxicity of PVA-DCA and PVA-UDCA against MiaPaCa-2 cells at pH = 7.4 (black) and 6.5 (red) after 24 h incubation at 37 °C. Cytotoxicity was evaluated by WST-8 assay. (b) Schematic illustration cytotoxicity assessment of PVA-UDCA pre-incubated or directly applied on MiaPaCa-2 cells. (c) Cell viability of MiaPaCa-2 cells treated with 0.01 mg mL^-1^ PVA-UDCA pre-incubated at 37 °C for 0 and 24 h at pH = 7.4 (black) and 6.5 (red). (d) Confocal microscopy images of MiaPaCa-2 cells after treatment with 0.1 mg mL^-1^ PVA-UDCA-TRITC at pH = 7.4 and 6.5 after 3 h incubation at 37 °C. Scale bars = 60 μm. (e) Number of PVA-UDCA-TRITC and PVA-DCA-TRITC aggregates per cell using MiaPaCa-2 cells treated with 0.1 mg mL^-1^ PVA-DCA-TRITC and PVA-UDCA-TRITC at pH = 7.4 (black) and 6.5 (red) at 3 h incubation at 37 °C. In (a), (c) and (e), n = 3 biologically independent samples. Statistical analysis was performed using unpaired two-tailed Student’s *t*-test. Data are presented as mean ± S.D. (\*\**p* < 0.01).

We revealed that the aggregate size of PVA-UDCA increased over time (Figure S13). Using this result, MiaPaCa-2 cells were treated with PVA-UDCA pre-incubated for 24 h and the cell viability was measured to investigate the contribution of aggregation to cytotoxicity (Figure 4b). The cell viability of MiaPaCa-2 cells treated with PVA-UDCA pre-incubated for 0 h at a concentration of 0.01 mg mL^-1^ was 93% and 90% at pH = 7.4 and 6.5, respectively, suggesting low cytotoxicity at both pHs (Figure 4c). In contrast, those pre-incubated for 24 h showed significant pH-responsive cytotoxicity at lower concentrations than those pre-incubated for 0 h, 111% at pH = 7.4 and 42% at 6.5, indicating the sufficient contribution of aggregation property of PVA-UDCA to cytotoxicity. These results suggested that pH-responsive cell adsorption of PVA-UDCA may contribute to the pH-responsive cytotoxicity.

To confirm our hypothesis, the adsorption of PVA-DCA and PVA-UDCA to the cell surface was investigated. PVA-DCA and PVA-UDCA were conjugated with fluorescent probes, tetramethyl rhodamine B isothiocyanate (PVA-DCA-TRITC and PVA-UDCA-TRITC), and cell adsorption of both polymers was evaluated by fluorescence imaging. Confocal images of MiaPaCa-2 treated with PVA-DCA-TRITC at a concentration of 0.1 mg mL^-1^ after 3 h of incubation showed PVA-DCA-TRITC aggregation and cell adsorption at pH = 7.4 and 6.5, suggesting no pH-responsive aggregation and cell adsorption (Figure S16). In contrast, those of PVA-UDCA-TRITC revealed aggregates number and cell adsorption markedly increased at pH = 6.5 compared to pH = 7.4, suggesting pH-responsive aggregation and cell adsorption of PVA-UDCA-TRITC (Figure 4d). The total area of aggregates of PVA-DCA-TRITC and PVA-UDCA-TRITC per cell were quantitatively evaluated using confocal images as shown in Figure S16 and Figure 4d, respectively. The number of PVA-DCA aggregates was 0.72 per cell at pH = 7.4 and 0.52 per cell at pH = 6.5 (Figure 4e). Those of PVA-UDCA-TRITC were 0.20 per cell at pH = 7.4 and 0.83 per cell at pH = 6.5, suggesting that PVA-UDCA aggregates were four times more abundant at pH = 7.4 than at pH = 6.5. These differences yielded a *p*-value of 0.06, indicating this result was in good agreement with the size evaluation of PVA-UDCA. However, the confocal images showed an overlap of aggregates of each polymer and cells at pH = 7.4 and 6.5 after 24 h incubation, suggesting no pH-responsive cell adsorption (Figure S17). Moreover, the total area of aggregates for both polymers showed no significant difference at pH = 7.4 and 6.5 (Figure S18). These results indicate that the pH-responsive cytotoxicity of PVA-UDCA after 24 h incubation is not due to pH-responsive cell adsorption.

The pH-responsive cytotoxicity of PVA-UDCA might be attributed to drastic hydrophobicity changes of UDCA in response to pH. The mechanism of the cytotoxicity of PVA-DCA and PVA-UDCA is depicted in Figure 5. PVA-DCA at pH = 7.4 showed self-aggregation in a few hours and adsorbed on cell surface, resulting in cell membrane disruption. After 24 h incubation, PVA-DCA formed microscale aggregates and showed higher cytotoxicity due to high hydrophobicity of DCA (Figure 5a). On the other hand, PVA-DCA at pH = 6.5 formed larger aggregates compared with that at pH = 7.4 after a few hours due to increasing hydrophobicity by protonation of DCA (Figure 5b). However, the cytotoxicity of PVA-DCA at pH = 6.5 was similar to that at pH = 7.4, suggesting that the DCA moiety is buried in the aggregate due to its high hydrophobicity. Conversely, PVA-UDCA at pH = 7.4 well dispersed at the nanoscale after several hours (Figure 5c). The polymer showed significantly lower cytotoxicity despite the formation of microscale aggregates after 24 hours of incubation, which may be attributed to the low hydrophobicity of UDCA. When UDCA was protonated at pH = 6.5, it became more hydrophobic and therefore aggregated and adsorbed on the cell surface after a few hours (Figure 5d). PVA-UDCA aggregates induced cell death more effectively than PVA-DCA aggregates after 24 h of incubation. This may be due to the distribution of UDCA moiety on the surface of the aggregates.

**Figure 5.**
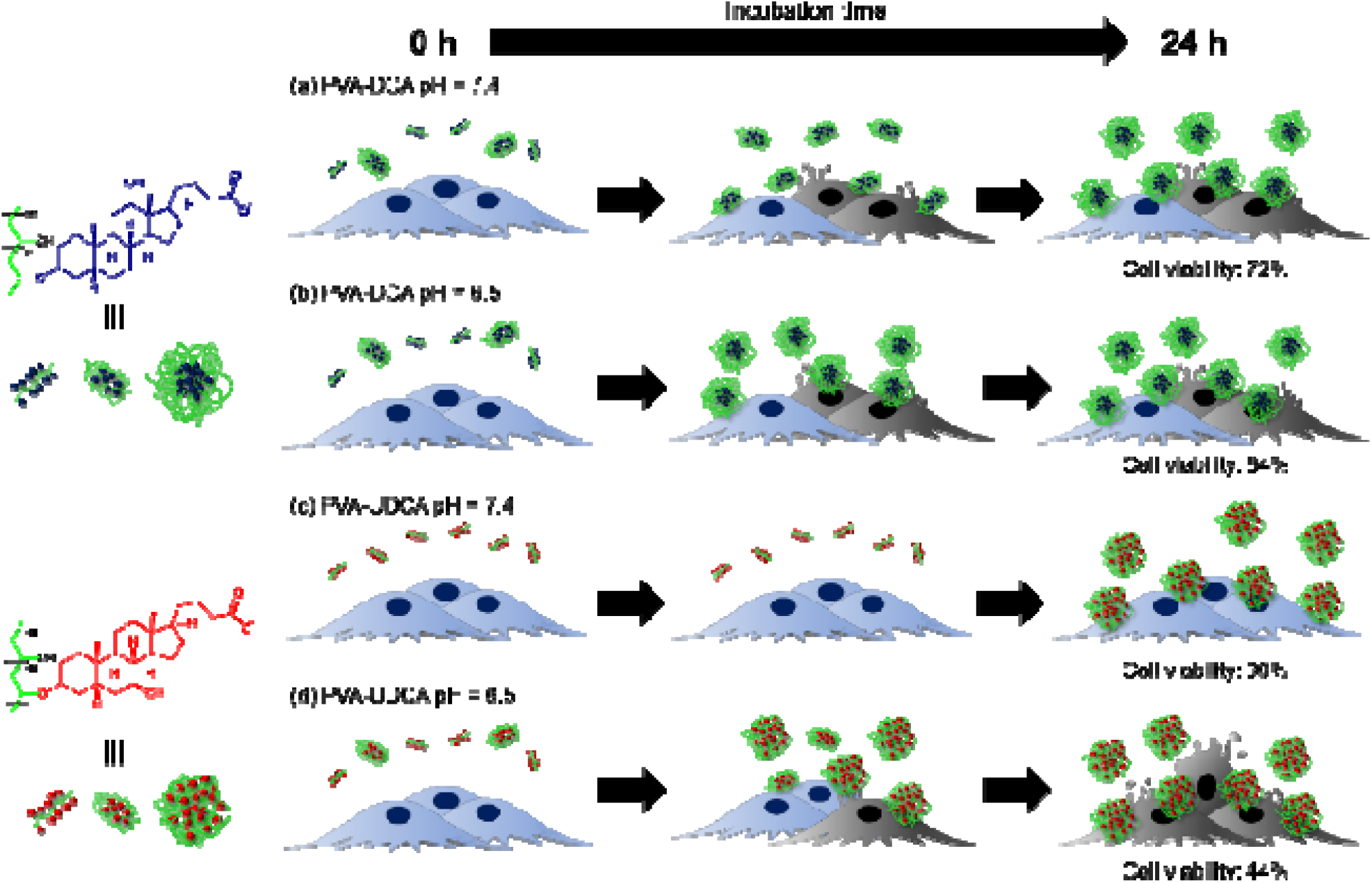
Schematic illustration of cancer cell death induction for PVA-DCA and PVA-UDCA over time. PVA-DCA formed aggregates at nano to micro scale over time and showed considerable cytotoxicity at pH = 7.4 due to its high hydrophobicity (a). Although PVA-DCA at pH = 6.5 formed aggregates after a few hours due to increasing hydrophobicity by protonation of DCA, cell membrane insertion is difficult due to the encapsulation of DCA in the core of the aggregate after 24 h incubation (b). However, PVA-UDCA at pH = 7.4 dispersed at the nanoscale aggregates and induced low cytotoxicity due to its low hydrophobicity (c), while it could be easy to form aggregates and insert to cell membrane at pH = 6.5 due to the protonation of UDCA on the outside of the aggregates (d).

## 4. CONCLUSION

In summary, we have developed a new class of MB, named PVA-UDCA, with high pH-responsive cytotoxicity by optimizing the bile acid moiety. PVA-UDCA achieved pH-responsive cytotoxicity derived from its low hydrophobicity at pH = 7.4 and high hydrophobicity due to protonation of UDCA. This study also revealed that a significant hydrophobic change at neutral pH and weakly acidic pH is important for the high pH-responsive aggregation property and cytotoxicity. Therefore, PVA-UDCA is expected to be applied as a novel cancer therapeutic molecule.

## ASSOCIATED CONTENT

### Supporting Information

The Supporting Information is available free of charge via the Internet at http://pubs.acs.org. Additional data analysis includes ^1^H NMR spectra, FT-IR spectra, CAC measurements, microscopy images, and cell viability.

## AUTHOR INFORMATION

### Corresponding author

*

### Author Contributions

K.M. carried out the experiments. K.M. wrote the manuscript. K.M., M.N., M.M. edited the manuscript. K.M., M.N., M.M. contributed to the design and implementation of the research and the analysis of results. All authors have seen and approved the final manuscript.

## ACKNOWLEDGEMENT

This work was supported by a Grant-in-Aid for Scientific Research (A) (20H00665), Bilateral Joint Research Projects of the JSPS (20199946), AMED-MPS (19be0304207h003), AMED-Translational Research Grant (H04), and in part by Japan Society for the Promotion of Science (JSPS) Grant-in-Aid for Young Scientists (Start-up) (JP20K22499) and JSPS Grant-in-Aid for Early-Career Scientists (JP22K18195). English proofreading was also supported by a Grant-in-Aid for Scientific Research (A) (20H00665).

---

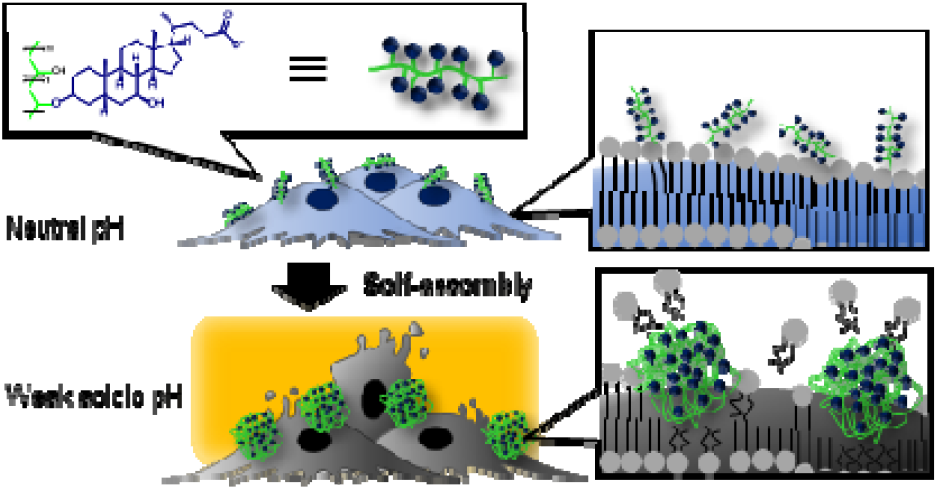
For table of contents only.

