## Supplementary information for "Fabrication of Molecular Blocks with High Responsiveness to the Cancer Microenvironment by Ursodeoxycholic Acid"

Corresponding Author:

Michiya Matsusaki, PhD., Professor

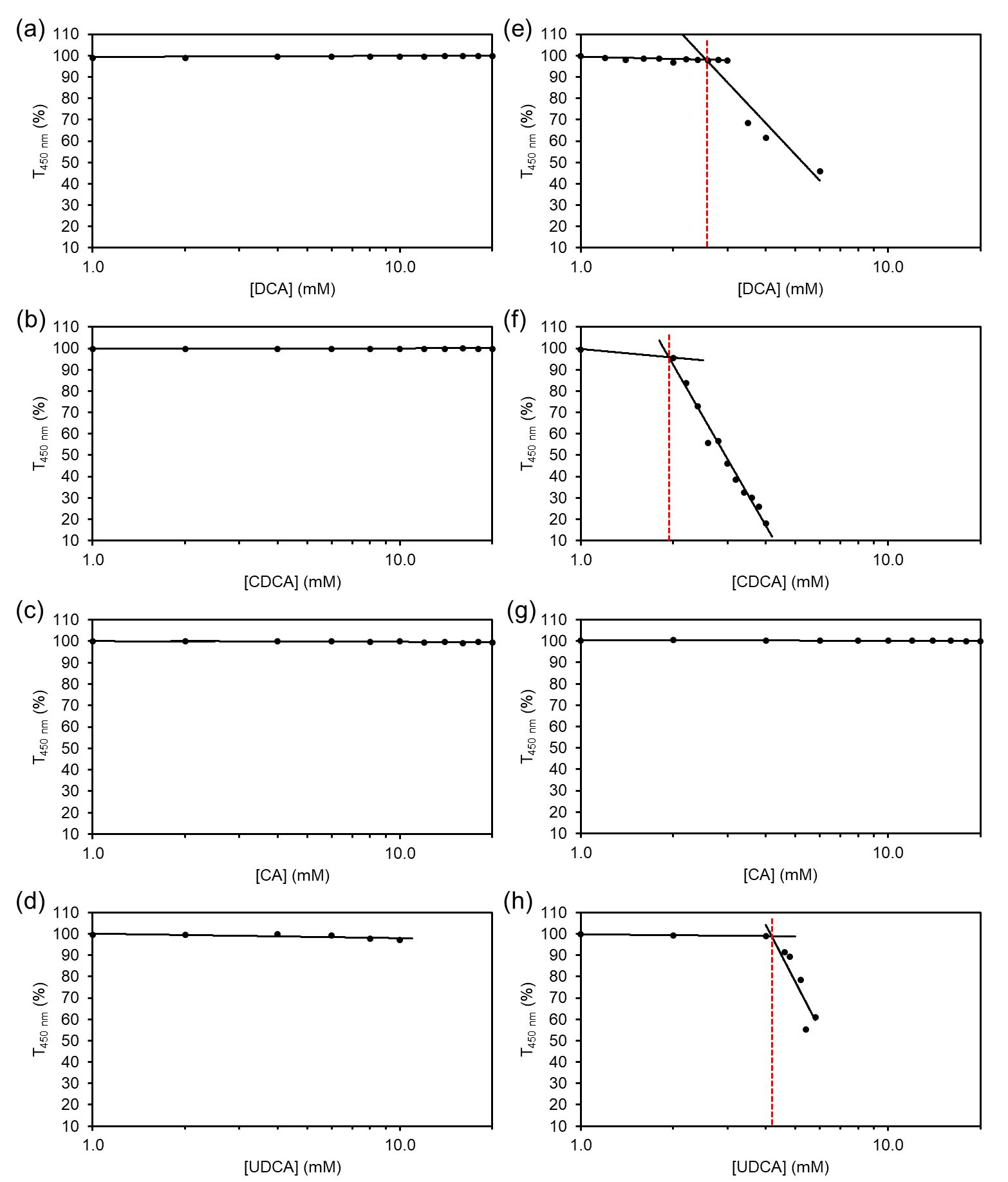

**Figure S1.** The critical aggregation concentration (CAC) evaluation by measurement of at wavelength of 450 nm (T_450 nm_). Concentration-dependent of T_450 nm_ of DCA at pH = 7.4 (a), 6.5 (e), CDCA at pH = 7.4 (b), 6.5 (f), CA at pH = 7.4 (c), 6.5 (g), UDCA at pH = 7.4 (d) and 6.5 (h) in phosphate-buffered saline (PBS) immediately after preparation at 37 °C.

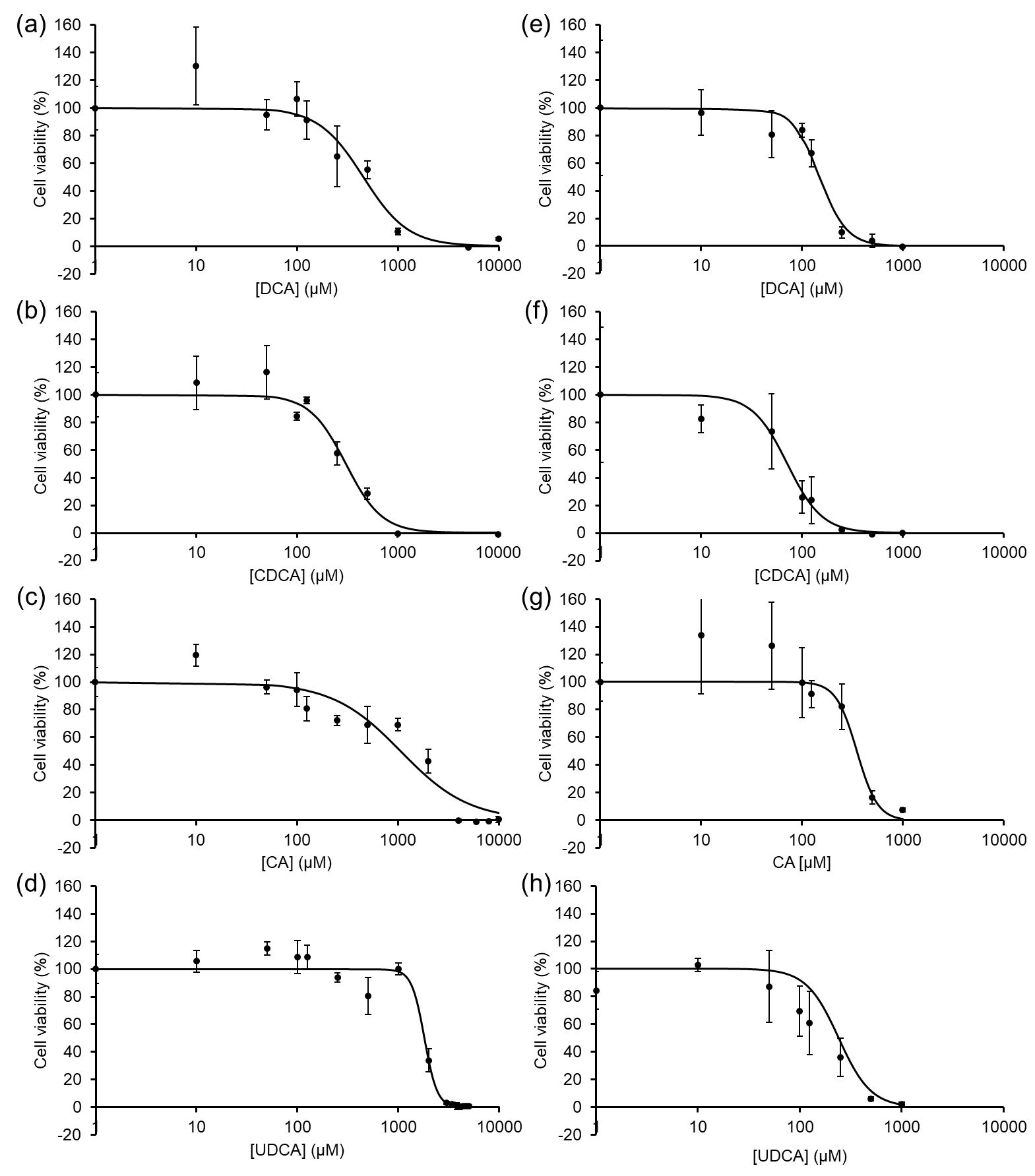

**Figure S2.** Concentration-dependent cytotoxicity of DCA at pH = 7.4 (a), 6.3 (e), CDCA at pH = 7.4 (b), 6.3 (f), CA at pH = 7.4 (c), 6.3 (g) UDCA at pH = 7.4 (d) and 6.3 (h) against MiaPaCa-2 cells after 24 h incubation at 37 °C. n = 3 biologically independent samples, and data are presented as mean ± S.D.

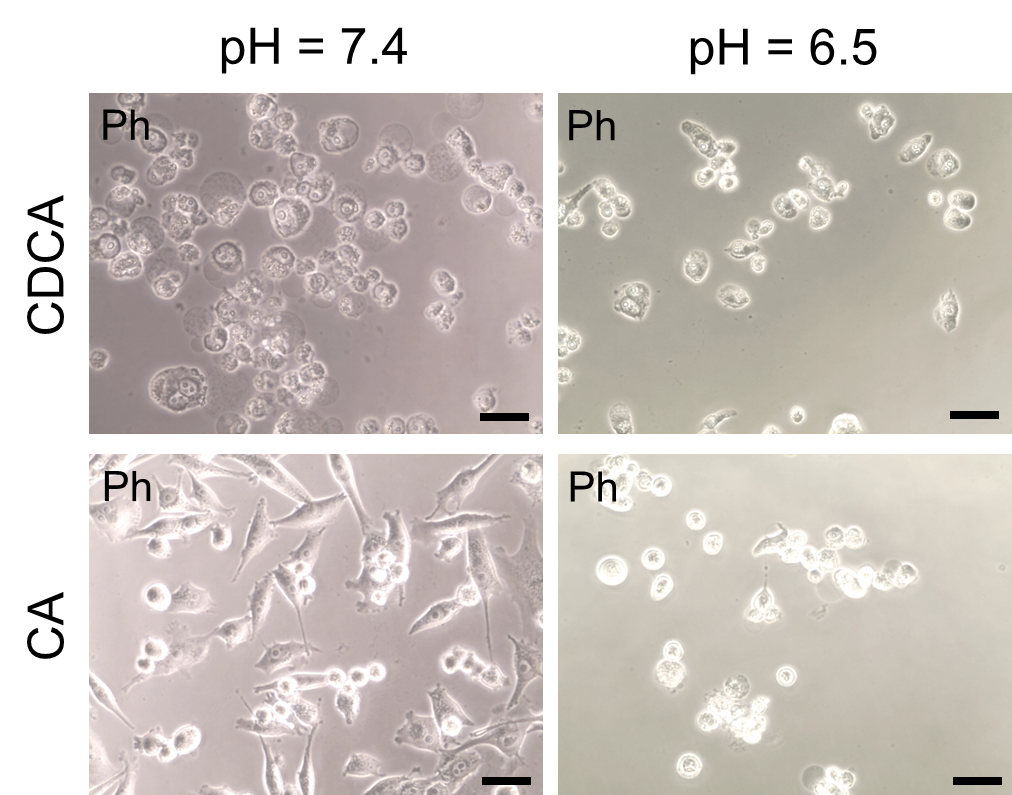

**Figure S3.** Phase contrast (Ph) microscopy images of MiaPaca-2 cells treated with 1000 µM CDCA and CA at pH = 7.4 and 6.3 for 24 h incubation at 37 ^o^C. Scale bars = 50 µm.

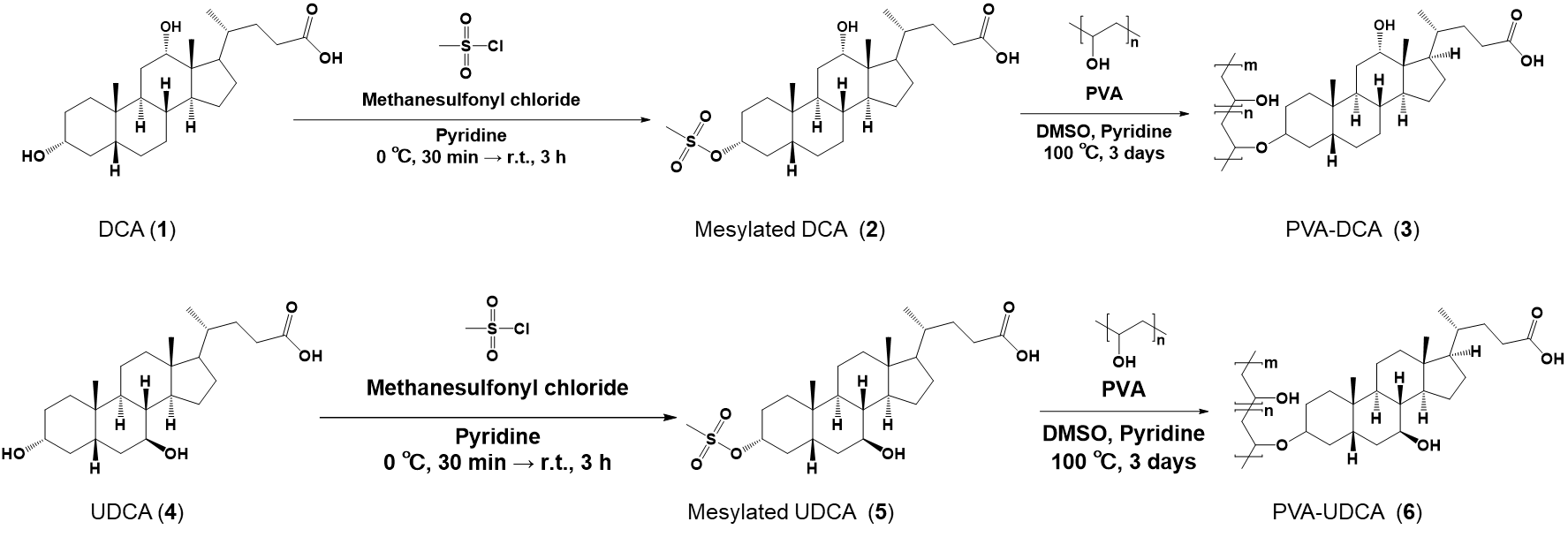

**Scheme S1.** Synthetic scheme of PVA-DCA and PVA-UDCA.

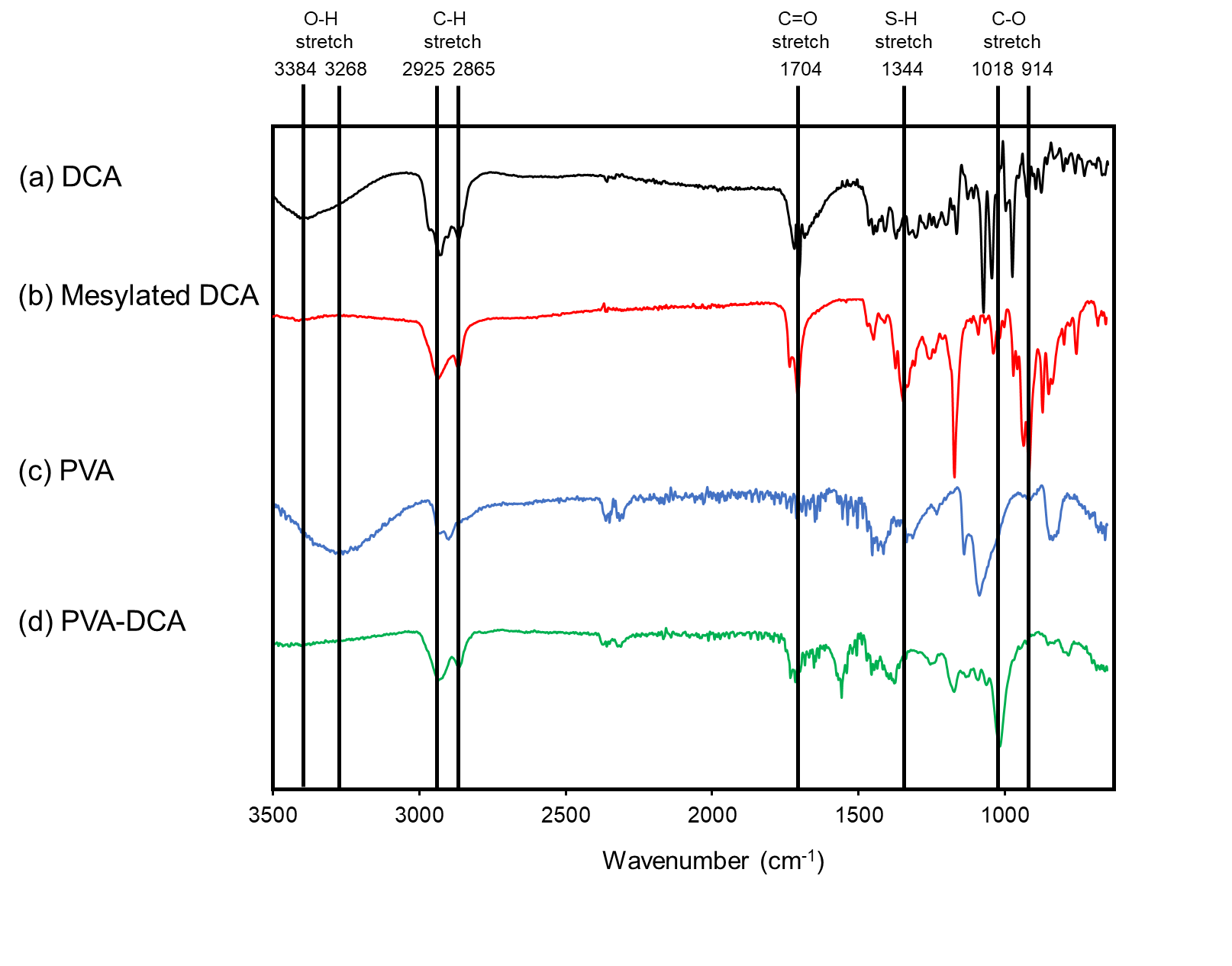

**Figure S4.** IR spectra of DCA (a), mesylated DCA (b), PVA (c) and PVA-DCA (d).

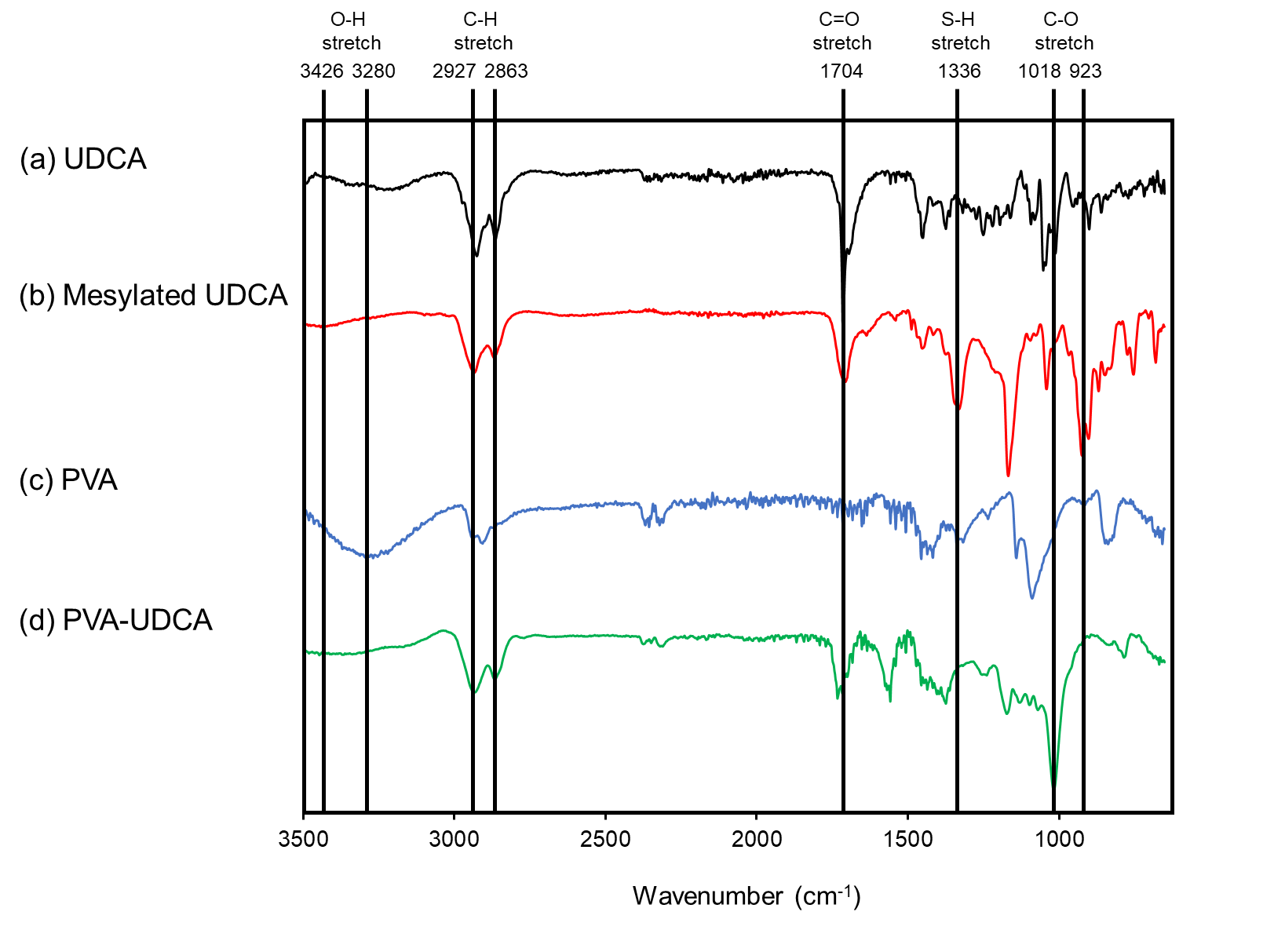

**Figure S5.** IR spectra of UDCA (a), mesylated UDCA (b), PVA (c) and PVA-UDCA (d).

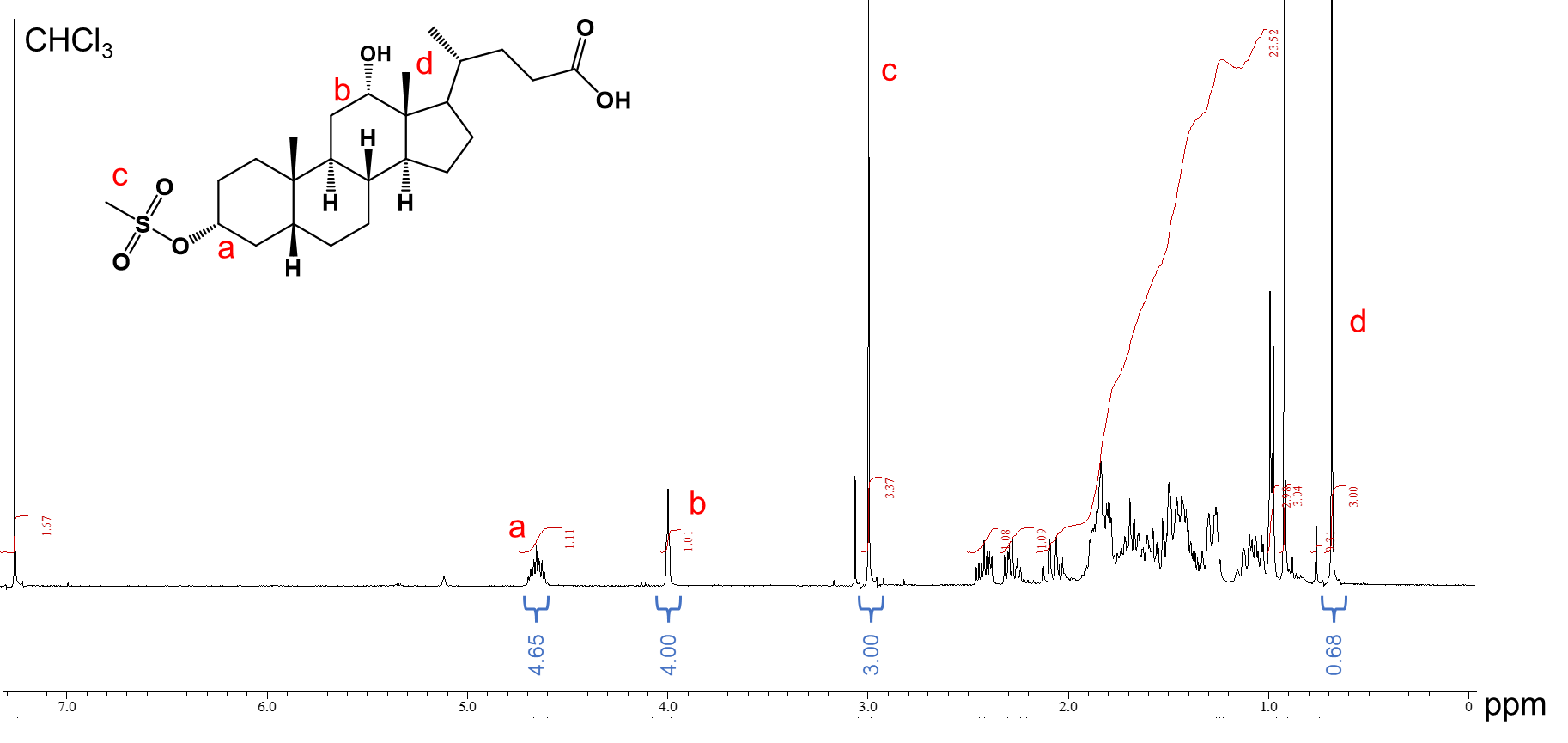

**Figure S6.**^1^H-NMR spectrum (Chloroform-*d*, 400 MHz, 25 °C) of mesylated DCA.

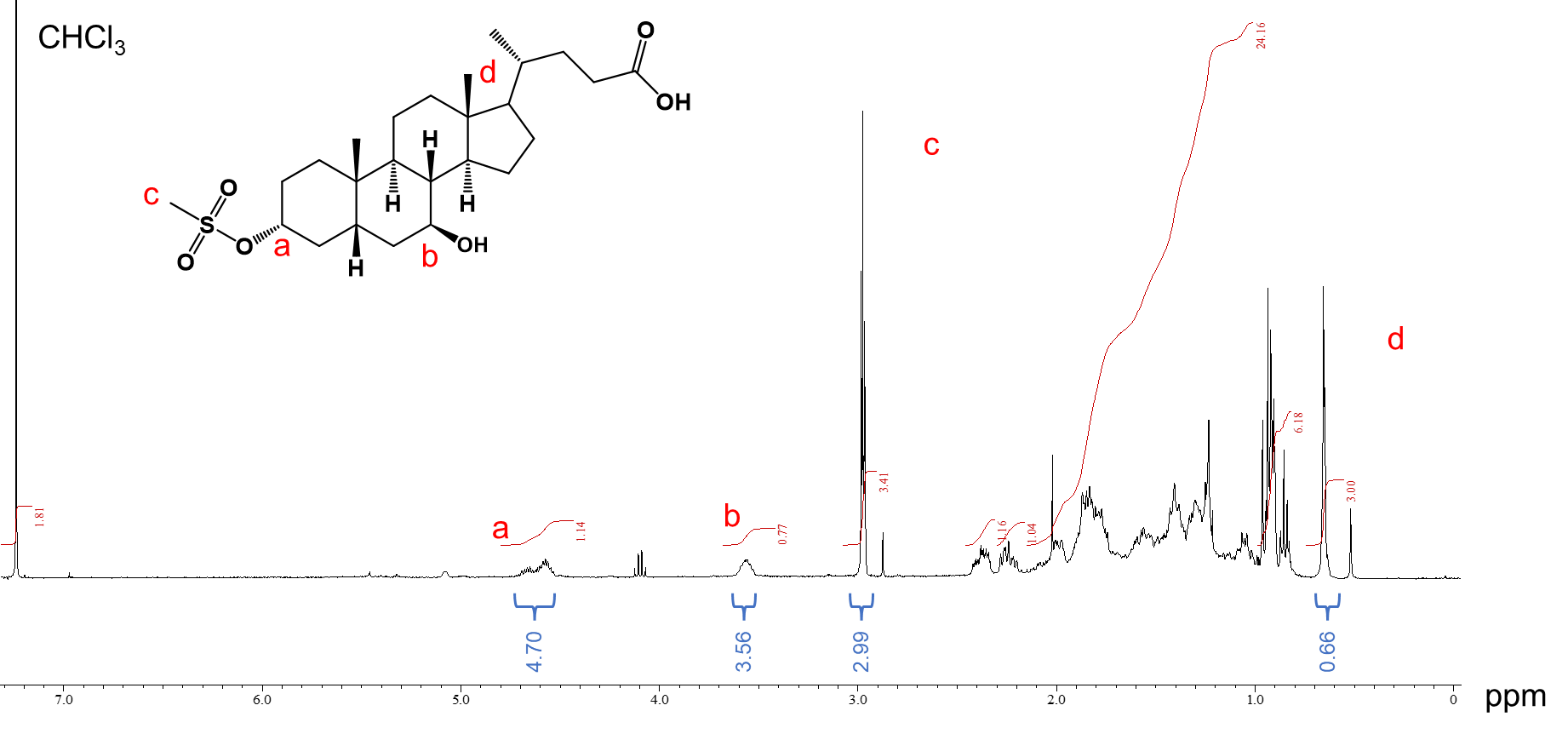

**Figure S7.** ^1^H-NMR spectrum (Chloroform-*d* 400 MHz, 25 °C) of mesylated UDCA.

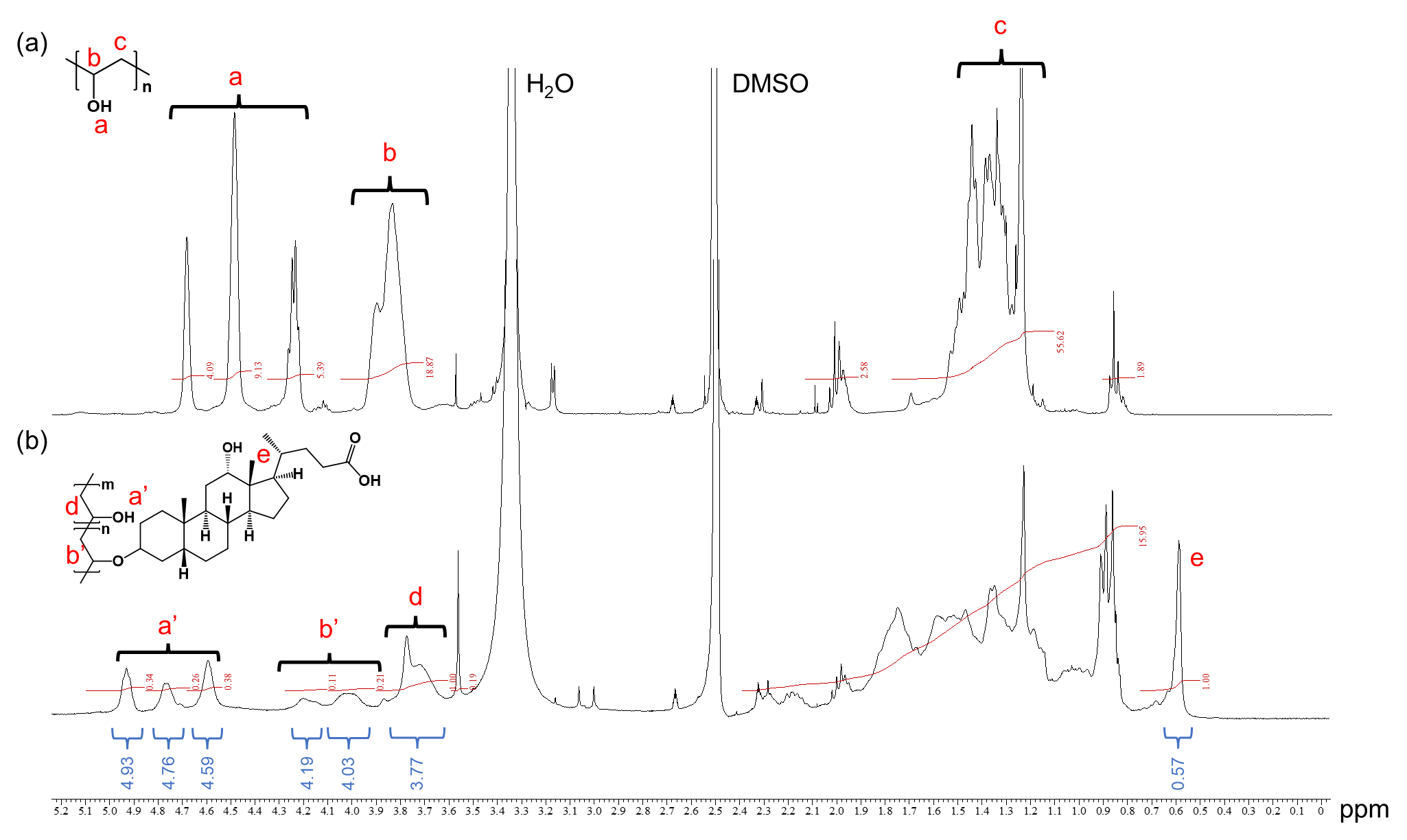

**Figure S8.** ^1^H-NMR spectra (DMSO-*d*_6_, 400 MHz, 25 °C) of PVA and PVA-DCA.

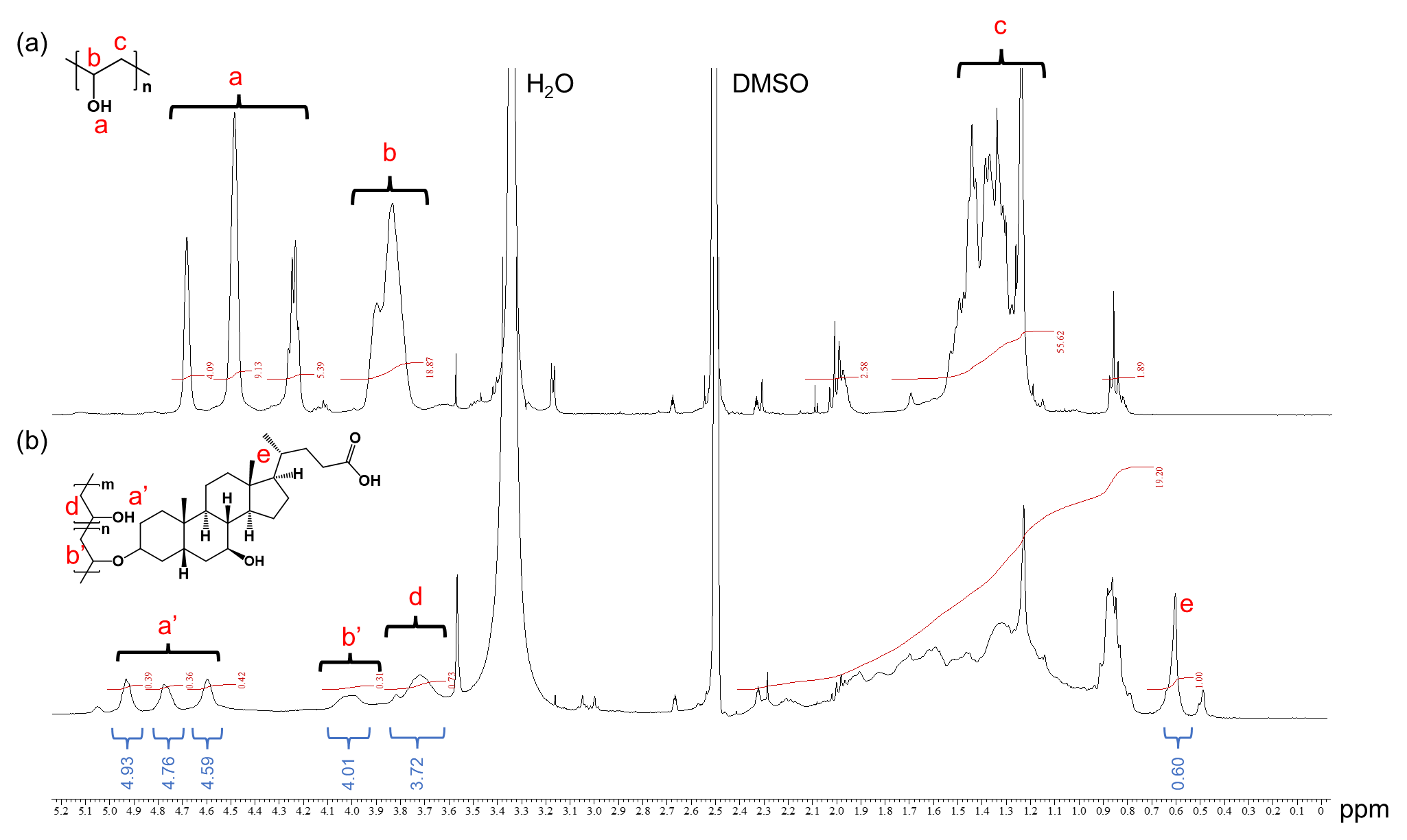

**Figure S9.** ^1^H NMR spectra (DMSO-*d*_6_, 400 MHz, 25 °C) of PVA and PVA-UDCA.

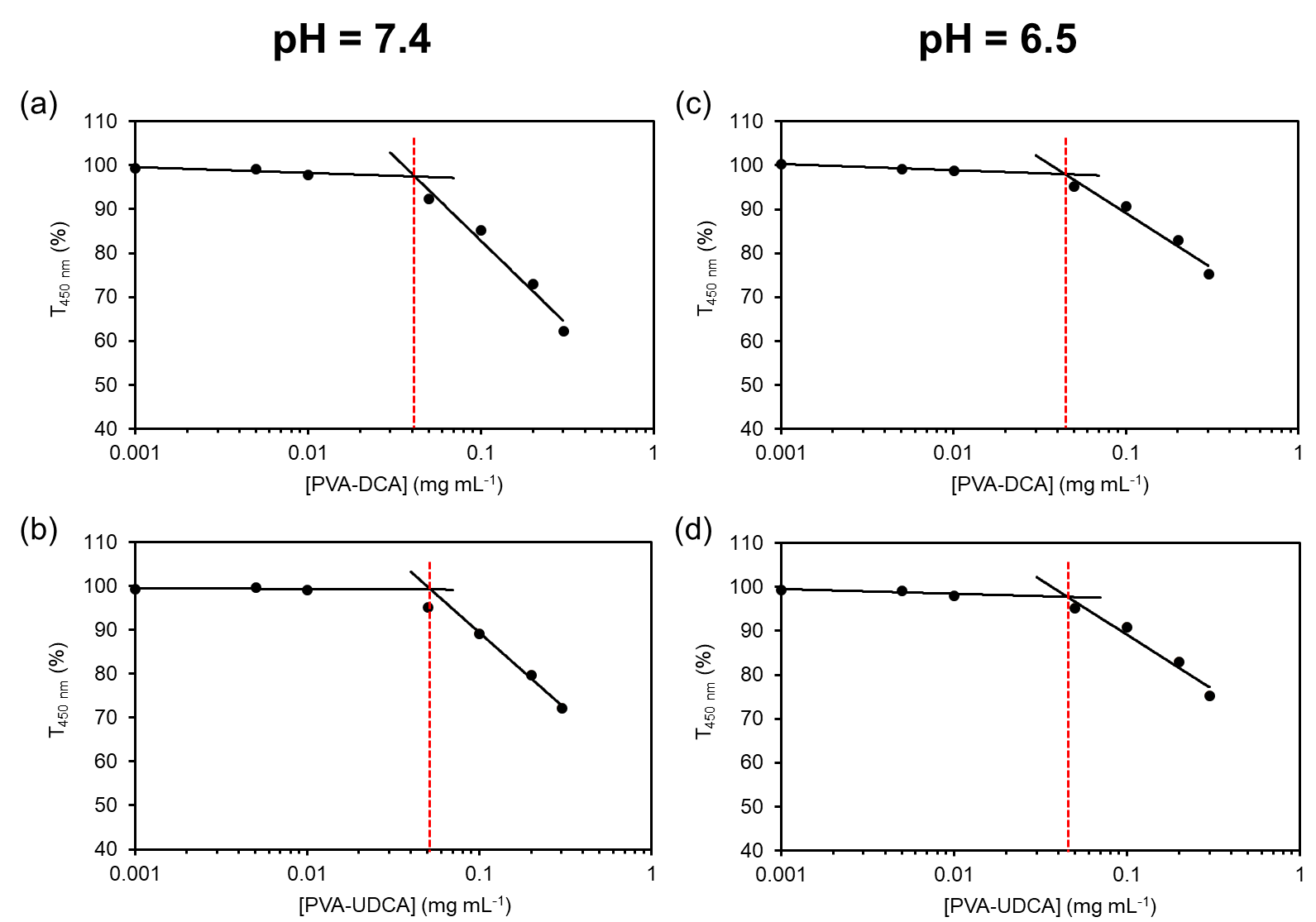

**Figure S10.** CAC evaluation by measurement of transmittance at a wavelength of 450 nm (T_450 nm_). Concentration-dependent of T_450 nm_ of PVA-DCA in PBS at pH = 7.4 (a), at pH = 6.5 (c). PVA-UDCA in PBS at pH = 7.4, (b) and PVA-UDCA in PBS at pH = 6.5 (d) immediately after preparation at 37 °C.

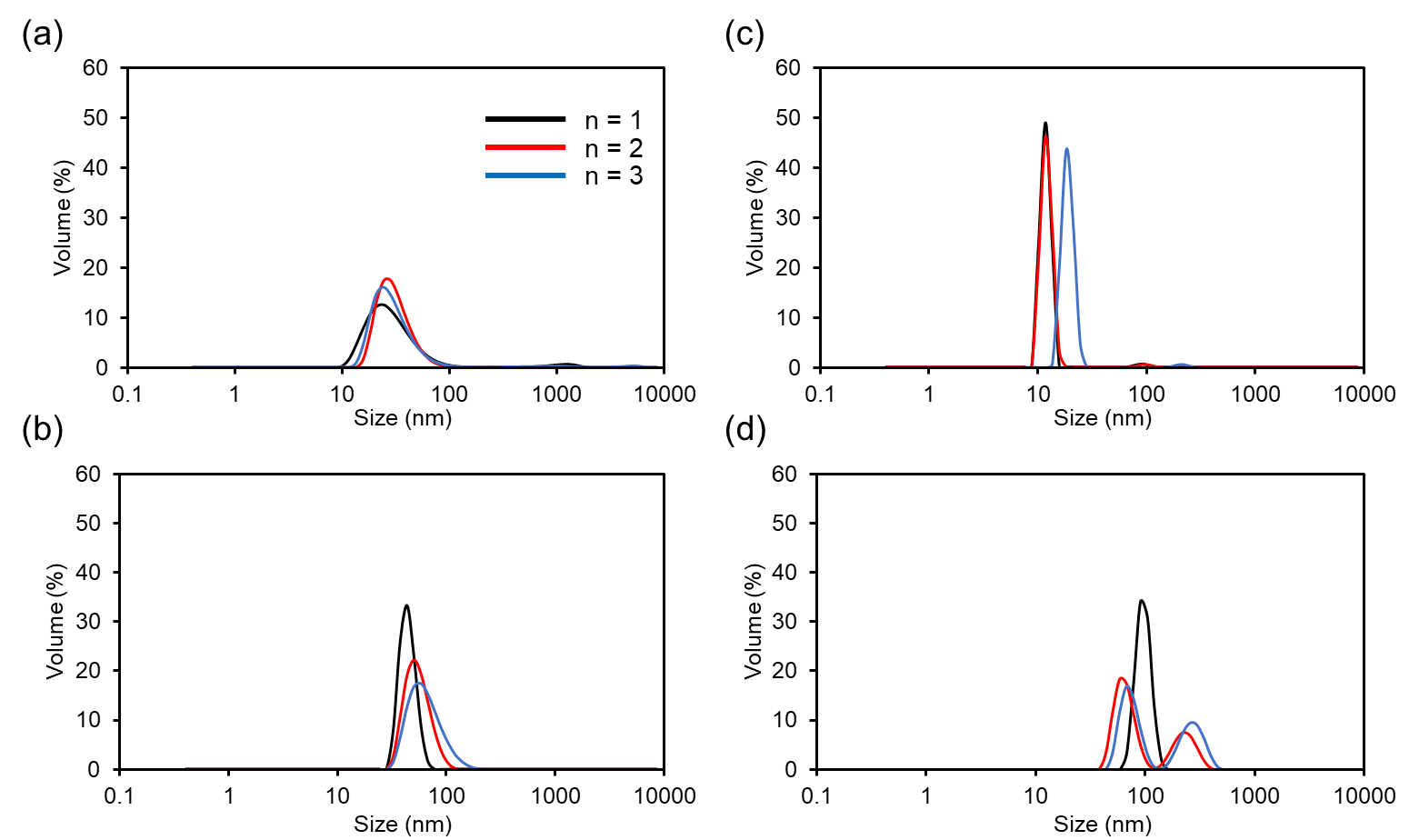

**Figure S11.** Size distribution of PVA-DCA in PBS at 0.1 mg mL^-1^ at pH = 7.4 (a) and pH = 6.5 (b), and PVA-UDCA in PBS at pH = 7.4 (c) and pH = 6.5 (d) at 37 °C immediately after preparation evaluated by dynamic light scattering (DLS).

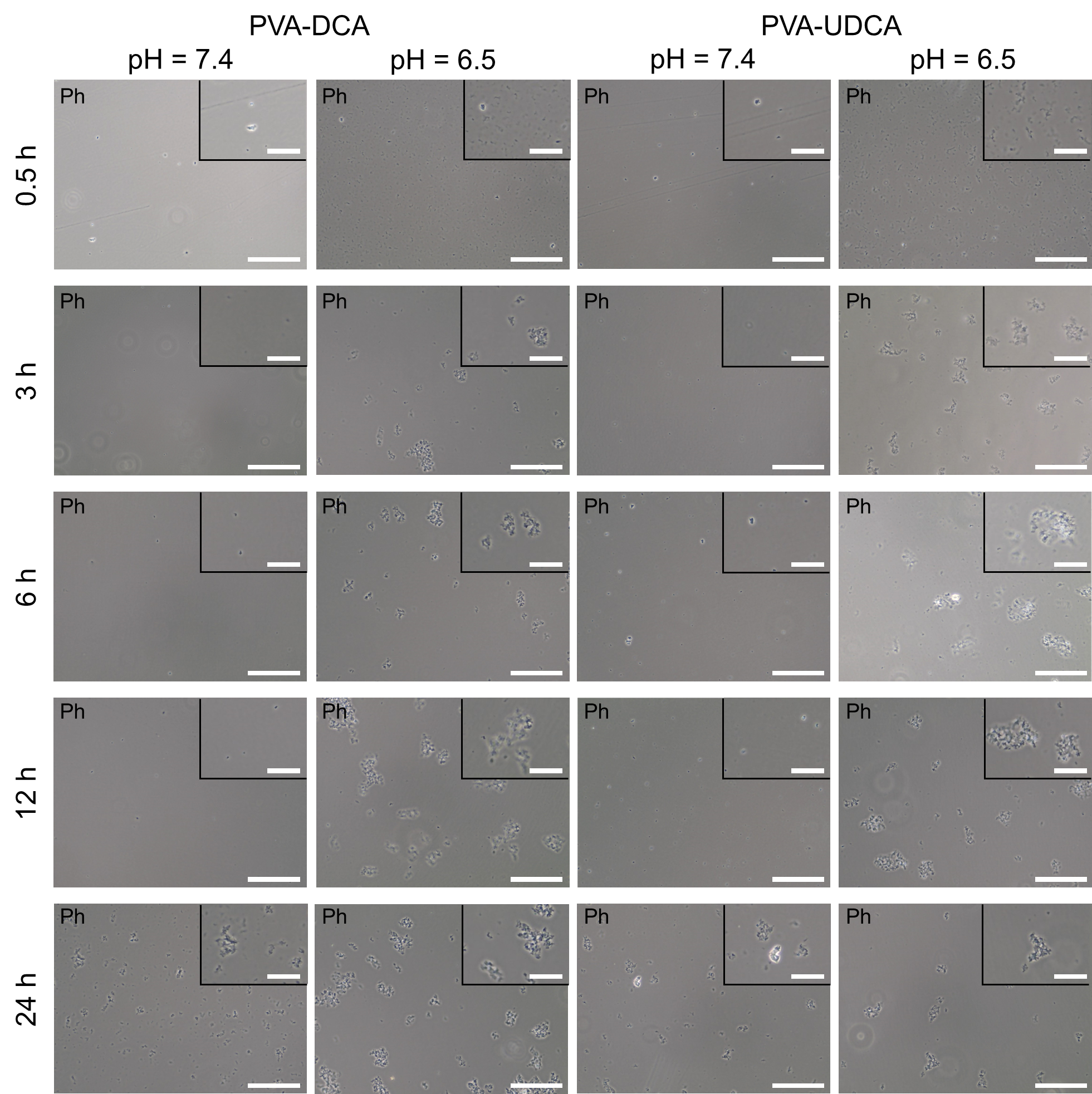

**Figure S12.** Phase contrast (Ph) microscopy images of 0.1 mg mL^-1^ PVA-DCA and PVA-UDCA solutions prepared in PBS at pH = 7.4 and pH = 6.5 after incubation 0.5, 3, 6, 12 and 24 h at 37 °C. Scale bars for images and enlarged images are respectively 50 µm and 20 µm.

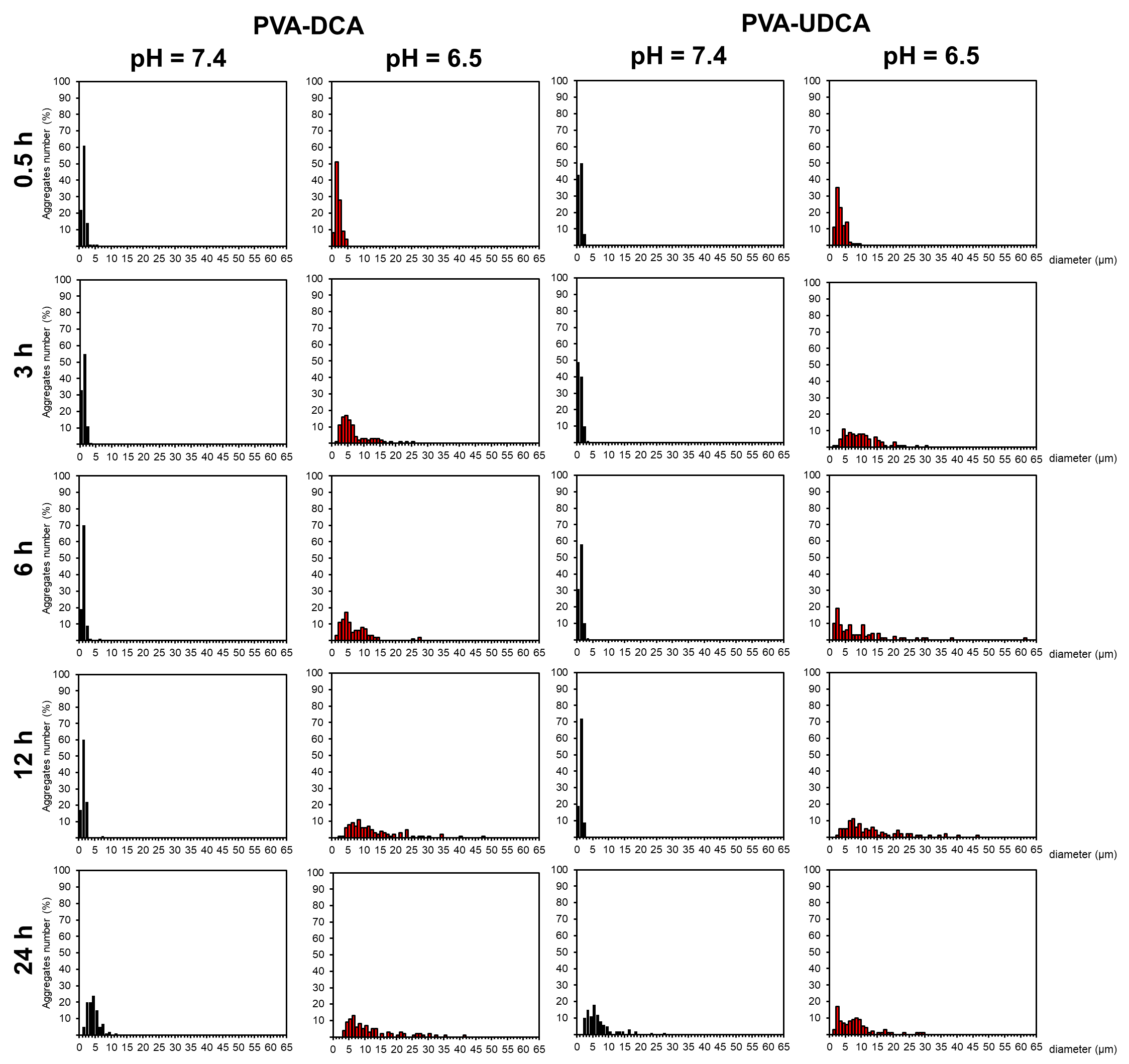

**Figure S13.** Size distribution of 0.1 mg mL^-1^ PVA-DCA and PVA-UDCA in PBS after incubation for 0.5, 3, 6, 12, and 24 h at pH = 7.4 (black) and pH = 6.5 (red).

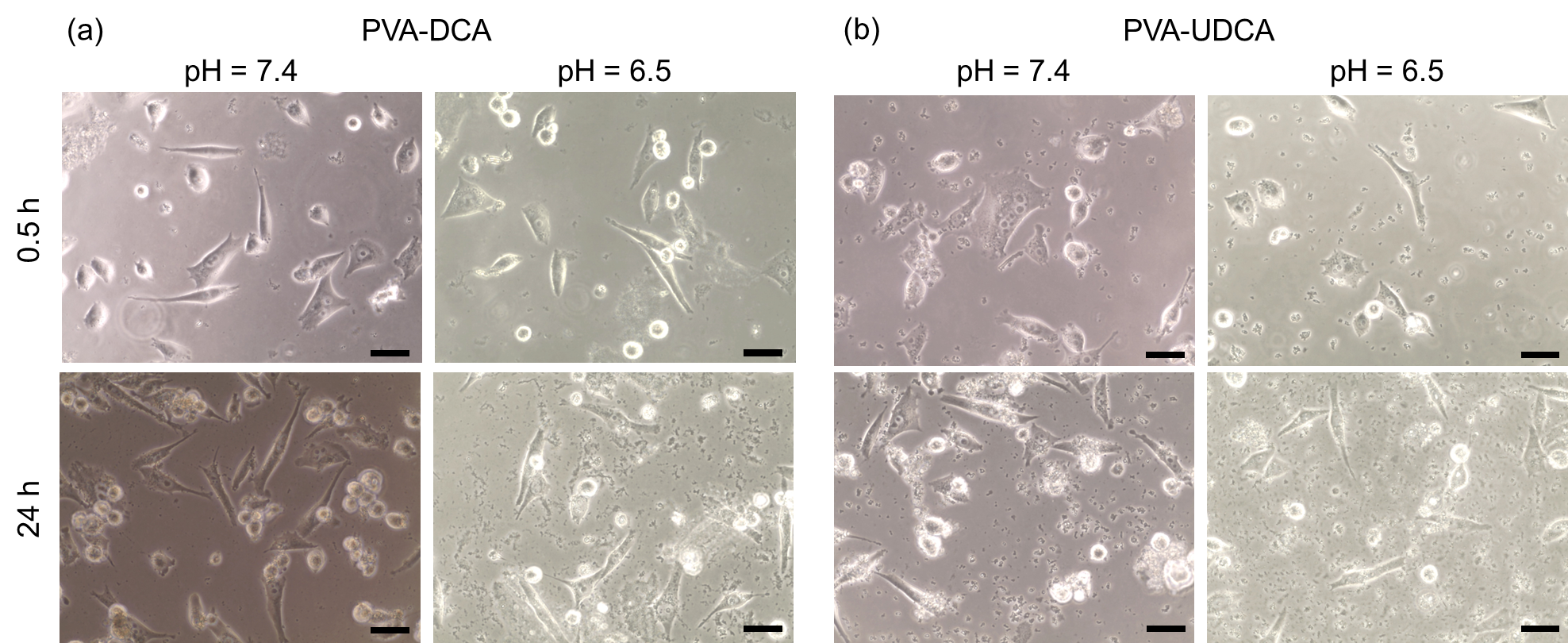

**Figure S14.** Phase contrast (Ph) microscopy images of MiaPaCa-2 cells treated with 0.1 mg mL^-1^ PVA-DCA (a) and PVA-UDCA (b) at pH = 7.4 and 6.5 for 0.5 h and 24 h incubation at 37 °C. Scale bars = 50 µm.

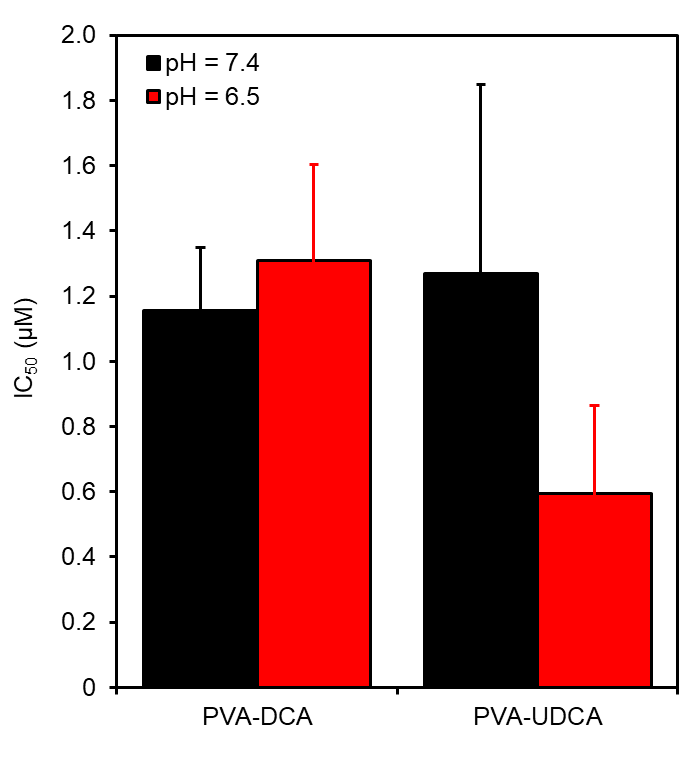

**Figure S15.** The IC_50_ values of PVA-DCA and PVA-UDCA using MiaPaCa-2 at pH = 7.4 (black) and 6.5 (red) after 24 h incubation. IC_50_ values were calculated from sigmoidal curve fitting.

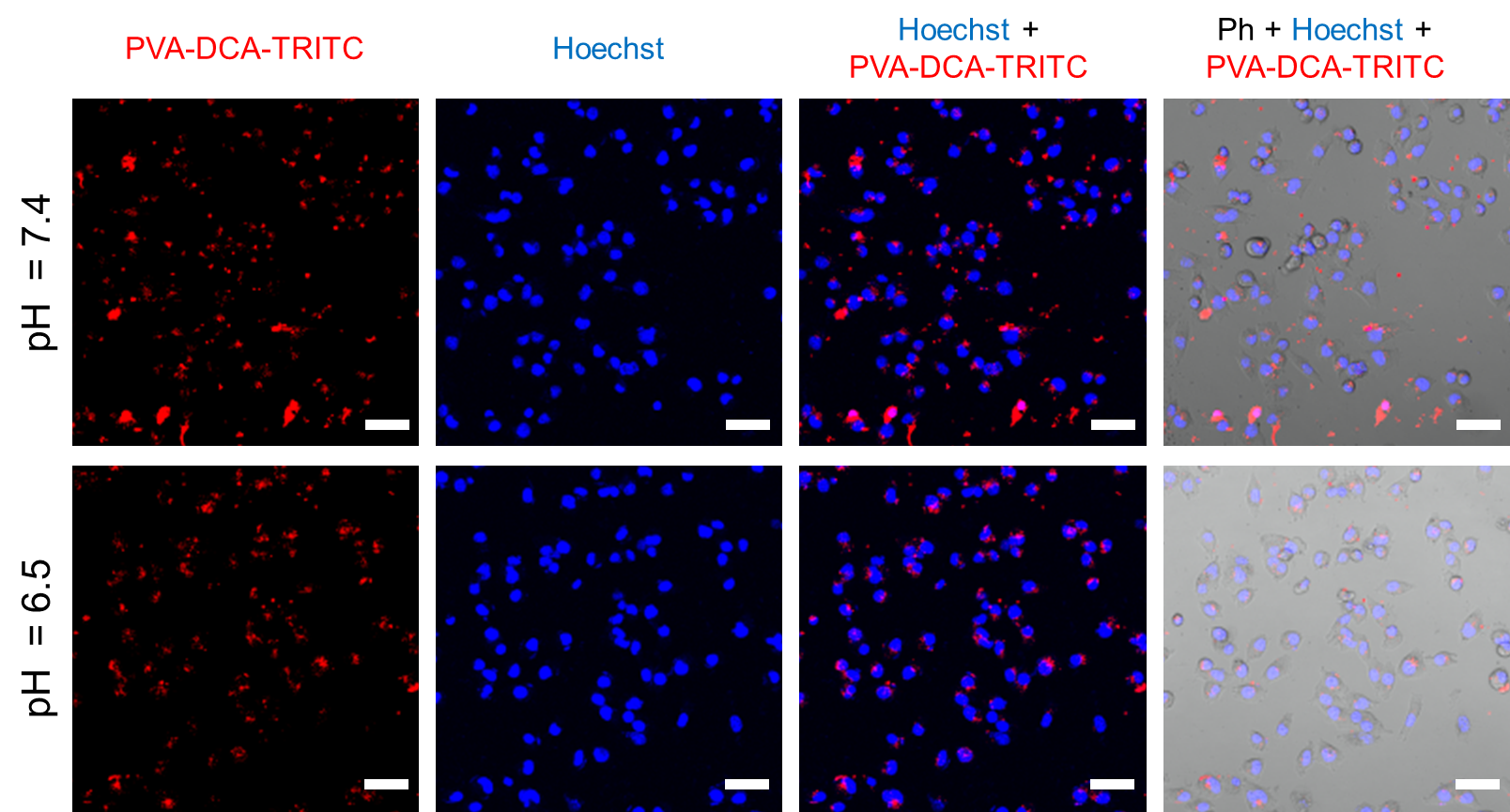

**Figure S16.** Confocal microscopy images of MiaPaCa-2 cells after 3 h incubation with 0.1 mg mL^-1^ PVA-DCA-TRITC at pH = 7.4 and 6.5 at 37 °C. Scale bars = 60 µm.

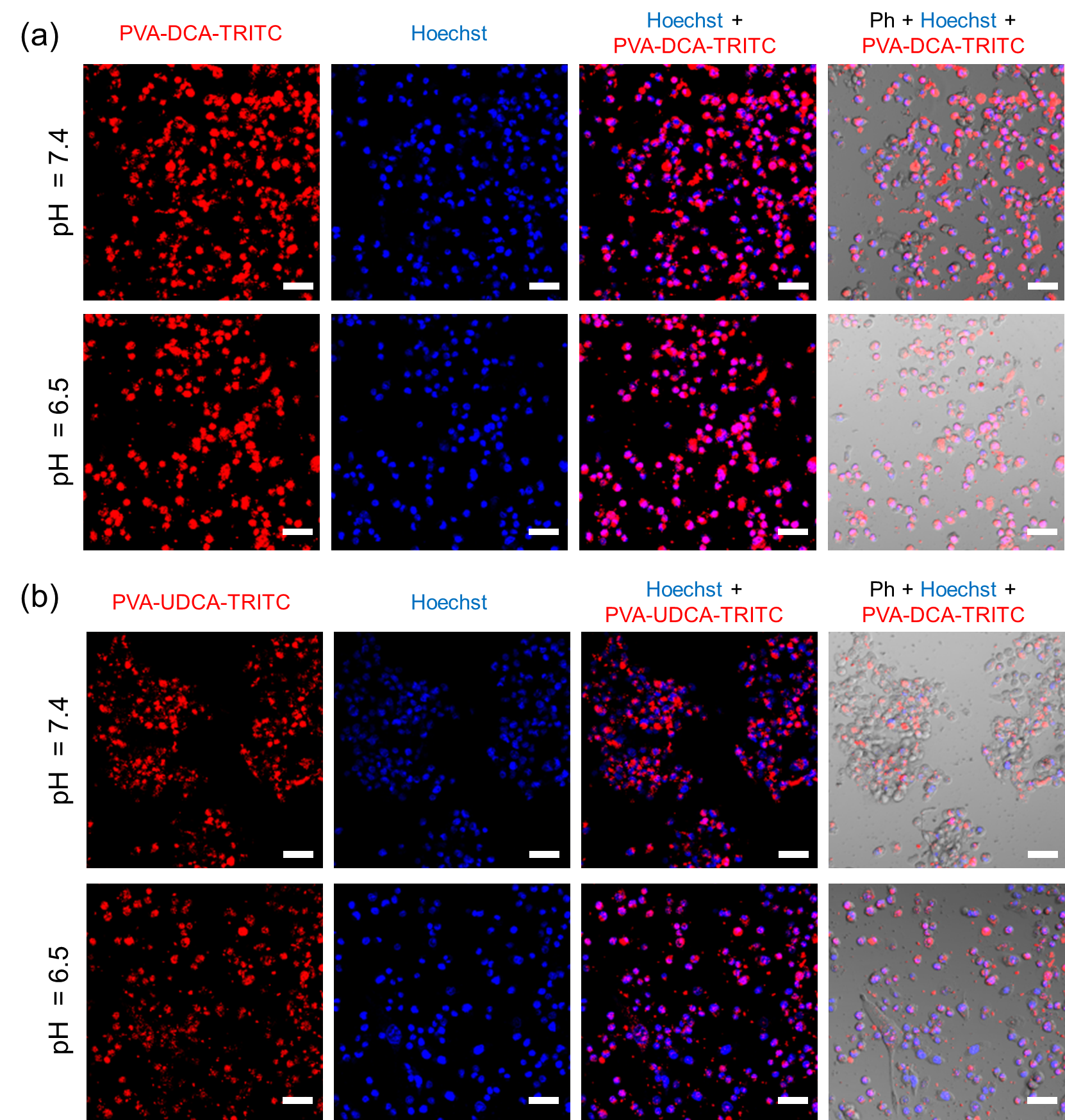

**Figure S17.** Confocal microscopy images of MiaPaCa-2 cells after 24 h incubation with 0.1 mg mL^-1^ PVA-DCA-TRITC (a) and PVA-UDCA-TRITC (b) at pH = 7.4 and 6.5 at 37 °C. Scale bars = 60 µm.

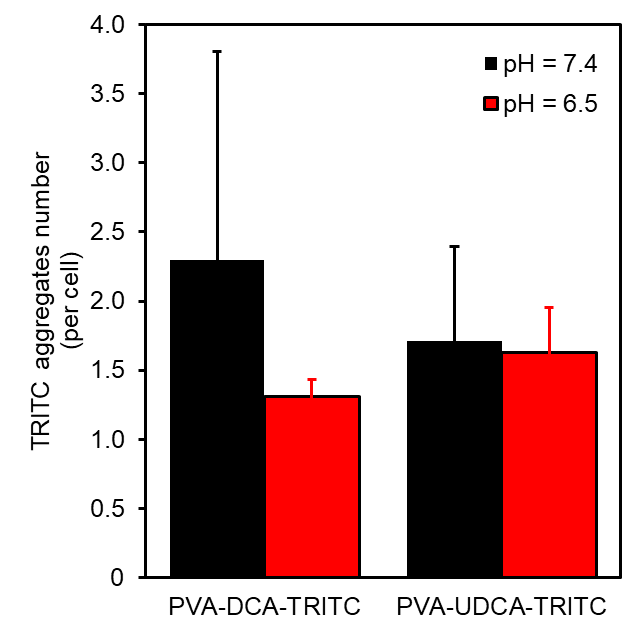

**Figure S18.** Number of TRITC aggregates per cell using MiaPaCa-2 cells after 24 h incubation with 0.1 mg mL^-1^ PVA-DCA and PVA-UDCA at pH = 7.4 (black) and 6.5 (red). n = 3 biologically independent samples. Data are presented as mean ± S.D.
